# Scalable TCR synthesis and screening enables antigen reactivity mapping in vitiligo

**DOI:** 10.1101/2025.04.24.649726

**Authors:** Stephanie A. Gaglione, Rachit S. Mukkamala, Chirag Krishna, Blake E. Smith, Marc H. Wadsworth, Scott A. Jelinsky, Caleb R. Perez, Laura Schmidt-Hong, Erica L. Katz, Kyle J Gellatly, Lestat R. Ali, Jiao Shen, Patrick V. Holec, Qingyang Henry Zhao, Amanda O. Chan, Ellen J. K. Xu, Kellie M. Kravarik, Julia A. Guzova, Connor S. Dobson, Harshabad Singh, Manuel Garber, Michael Dougan, Stephanie K. Dougan, John E. Harris, Aaron Winkler, Michael E. Birnbaum

## Abstract

T cell receptors (TCRs) mediate antigen recognition in adaptive immunity, yet large-scale mapping of TCR-antigen interactions remains a major challenge. Current approaches to synthesize and functionally screen TCRs remain technically complex and limited in throughput. We introduce a modular strategy, TCRAFT, to rapidly construct tens of thousands of TCRs for <$1 each while maintaining TCRα-β pairing with >99% accuracy. We integrate this approach with a high-throughput antigen discovery platform to enable library-on-library TCR-antigen screening. We reconstruct and screen 3,808 TCRs from vitiligo lesions, linking TCR specificity to transcriptional phenotypes for antigen-reactive T cells. To demonstrate scalability, we synthesize and screen 30,810 TCRs from donors with pancreatic ductal adenocarcinoma to capture antigen-specific TCRs. This workflow reduces the cost and complexity of large-scale TCR screening, enabling the expansion of the known landscape of antigen-specific TCRs in vitiligo with a method that can be readily extended to other immunological applications.

## Introduction

T cells are central to adaptive immunity, responding to specific antigens in the context of cancer, infection, and autoimmunity via T cell receptors (TCRs). Each T cell clone expresses a unique TCR, a heterodimer composed of a TCRα and TCRβ chain, that binds to short peptides presented by major histocompatibility complex (MHC) proteins (pMHCs) to elicit an antigen-specific T cell response. Screening TCRs for antigen reactivity at a high throughput would advance our understanding of natural immune function while providing sources of potentially therapeutically relevant TCRs and data for improving computational models of TCR specificity^1,2^. However, the vast diversity of the TCR repertoire—up to 10⁸ unique clonotypes per individual^3,4^—makes large-scale mapping of TCR-antigen interactions a major challenge.

Identifying antigen-specific TCRs and their associated cell phenotypes is a critical priority in cancer^5^ and autoimmune research^6^. T cells recognizing cancer-specific neoantigens often exhibit exhausted phenotypes^7,8^, and their role in tumor control is well-documented^9,10^. In contrast, the phenotypic states and antigenic targets of T cells driving autoimmunity are less well understood. Proposed sources of autoantigens include self-antigens^11^, post-translationally modified epitopes^12^, and viral epitopes^13^. Vitiligo is one of the few autoimmune diseases with known antigens, largely due to insights from melanoma research. In both diseases, autoreactive CD8^+^ T cells target melanocyte-derived antigens including MART-1, tyrosinase, and gp100^7,14–17^. However, the extent of overlap in TCR identity and transcriptional features of antigen-specific T cells between vitiligo and melanoma remains unclear. Defining shared and distinct features of these T cell populations is essential for advancing targeted therapies in both cancer^5^ and autoimmunity^6^.

Despite advances in single-cell and computational methods for sequencing and pairing TCR chains^18,19^, TCR datasets alone provide little insight into antigen specificity. Current screening strategies evaluate only a small subset of TCRs, typically selected based on clonotypic expansion or phenotypic markers^1^. Furthermore, assembling and screening TCR libraries remains costly, labor-intensive, and inefficient. Existing approaches rely on synthesizing expensive gene fragments, assembling individual TCRs in array using custom oligonucleotides and libraries of TCR variable regions^20–22^, or amplifying TCRs from primary cell samples^23–25^. A key limitation of synthetic TCR assembly is maintaining the natural pairing between TCRα and TCRβ chains, which is essential for preserving native TCR function. Current pooled TCR assembly methods either require co-transduction of separate TCR chains, an approach limited in scale, or scramble TCRα/β pairings, resulting in libraries dominated by non-functional TCRs^26^. While recent methods enable pooled assembly of up to 1,000 TCRs, error rates and technical complexity limit broad adoption or further increases in throughput^27^.

To address these limitations, we developed **TCR** Rapid **A**ssembly for **F**unctional **T**esting (TCRAFT), a scalable and cost-effective approach to easily synthesize tens of thousands of TCRs in a pool for under $1 per TCR. TCRAFT links pools of TCR variable region genes with CDR3α/β-containing oligonucleotides via three Golden Gate assembly steps, yielding paired-chain TCR libraries with >99% correctly assembled TCRs. We integrate TCRAFT with RAPTR^28^, a high-throughput antigen discovery platform, to enable one-pot library-versus-library screening of TCR reactivity. Using this system, we screened 3,808 TCR clonotypes from vitiligo lesions against 101 vitiligo and melanoma-associated antigens, extracting precise TCR-antigen pairings in a single step. We orthogonally validated these hits and expanded reactivity screening to 561 antigens using an antigen presentation assay. By linking TCR specificity to corresponding gene expression data, we identified correlations between transcriptomic signatures of vitiligo- and melanoma-associated antigen-reactive T cells. To further demonstrate scalability, we used TCRAFT to synthesize a single library of 30,810 TCRs from surgically resected tumors and PBMCs of patients with pancreatic ductal adenocarcinoma and identified antigen-specific TCRs using antigen-presenting cells pulsed with peptides.

By significantly reducing cost, labor, and technical barriers, TCRAFT provides an accessible and efficient strategy for assembling large TCR libraries. We couple these libraries with high-throughput antigen screening and single-cell transcriptomics to link TCR specificity to cell phenotype. Applied to vitiligo, this integrated approach uncovers transcriptional programs of melanocyte-reactive T cells. Together, these tools will accelerate efforts to decode TCR specificity, understand immune responses, and develop TCR-based immunotherapies.

## Results

### Modular assembly of pooled TCR libraries by hierarchical Golden Gate assembly

Single-cell and repertoire TCR sequencing enable deep profiling of clonal T cell dynamics in autoimmune diseases such as vitiligo, but TCR sequences provide limited disease context without antigen specificity. To address this, we aimed to establish a complete TCR-antigen discovery pipeline consisting of single-cell sequencing to extract paired TCR chain sequences, pooled synthetic TCR assembly, and high-throughput library-on-library TCR-antigen screening (**Figure 1A**). We first sought to address the bottleneck of assembling large TCR libraries from sequencing data.

**Figure 1.**
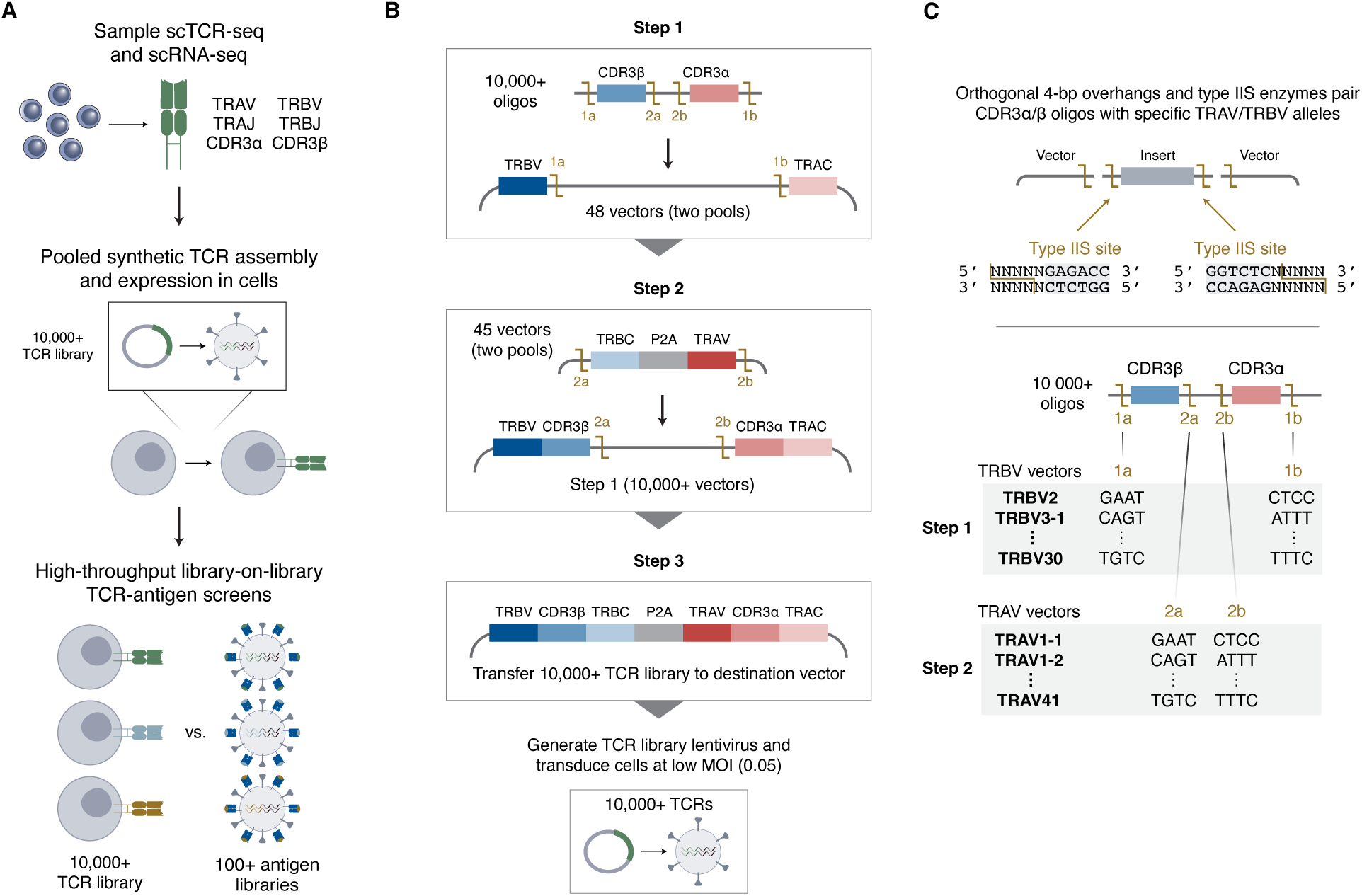
Overview of modular TCR assembly and screening pipeline. **(A)** Schematic of high-throughput screening pipeline to identify antigen-specific TCRs by sequencing samples to extract paired TCRαβ receptor sequences (step 1), assembling a TCR library in pool and expressing TCRs in cells (step 2), and pairing TCRs to cognate antigens in a library-on-library screen with lentiviruses pseudotyped with pMHCs or other antigen discovery methods (step 3). **(B)** Schematic of pooled TCR assembly method. Step 1 inserts oligos of paired CDR3β-α sequences into TRBV-TRAC vectors to generate TRBV-CDR3β-CDR3α-TRAC. Step 2 inserts TRBC-P2A-TRAV fragments between CDR3β and CDR3α to generate complete TCRs. Step 3 transfers the complete TCRs to an expression vector. TCR libraries are expressed in cells via lentiviral transduction at a low MOI. All steps are completed via Golden Gate assembly. **(C)** Schematic of approach to correctly pair TCR components. Type IIS enzymes generate 4-bp overhangs to facilitate ligation between reaction components. Each TRBV- and TRAV-containing vector contains unique pairs of 4-bp overhangs designed to maximize correct pairing while preserving coding sequences. Each CDR3α/β oligo contains four enzyme cut sites, with the outer 4-bp overhangs facilitating pairing to TRBV vectors and inner 4-bp overhangs corresponding to TRAV vectors.

To enable low-cost pooled assembly of TCRs from sequences, we generate libraries of linked TCRαβ receptors from pools of germline TRBV and TRAV vectors and oligonucleotide pools of linked CDR3α/β sequences (**Figure S1**). We first insert the CDR3α/β-containing oligos into vectors containing paired TRBV-TRAC sequences, followed by insertion of TRBC-P2A-TRAV sequences between the CDR3α and CDR3β sequences. The resulting product is a complete, synthetically formatted TCR (TRBV-TRBC-P2A-TRAV-TRAC), which is then transferred to an expression vector of choice via a third reaction (**Figure 1B**).

Our approach uses Golden Gate assembly to pair CDR3α/β fragments with TRAV and TRBV genes in a pooled setting (**Figure 1C**). This approach employs type IIS enzymes to generate four-base-pair overhangs, ensuring seamless linkage of coding regions. Although Golden Gate assembly has previously been demonstrated for assembling individual TCRs^22^, extending this approach to pooled library assembly remained infeasible due to the low orthogonality between four-base overhangs. To address this challenge, we leveraged a comprehensive 4-bp ligation fidelity dataset^29^ to design highly orthogonal pairs of overhangs for each vector in the pool (**Figure S2**). Each TRBV-TRAC and TRBC-TRAV fragment was assigned a distinct pair of 5’ and 3’ overhangs to facilitate precise assembly while preserving coding sequences (**Table S1**).

### Pooled assembly and characterization of a synthetic vitiligo 3,808 TCR library

To examine TCR reactivity in the context of vitiligo, we identified 3,808 TCRs in suction blister fluid from vitiligo lesions of 10 HLA-A*02:01^+^ (henceforth HLA-A2) donors (**Figure 2A, Table S2)**. While prior work suggests that vitiligo is driven by CD8^+^ T cells reactive to MART-1 and other melanocyte-specific antigens^30–33^, perilesional T cell reactivity has not been broadly examined due to limitations in screening T cells from tissue samples. Vitiligo offers a compelling context for studying antigen-specific T cell responses in autoimmunity given its strong HLA-A2 risk association^33,34^ and limited characterization of cellular mechanisms underlying a loss of tolerance.

**Figure 2.**
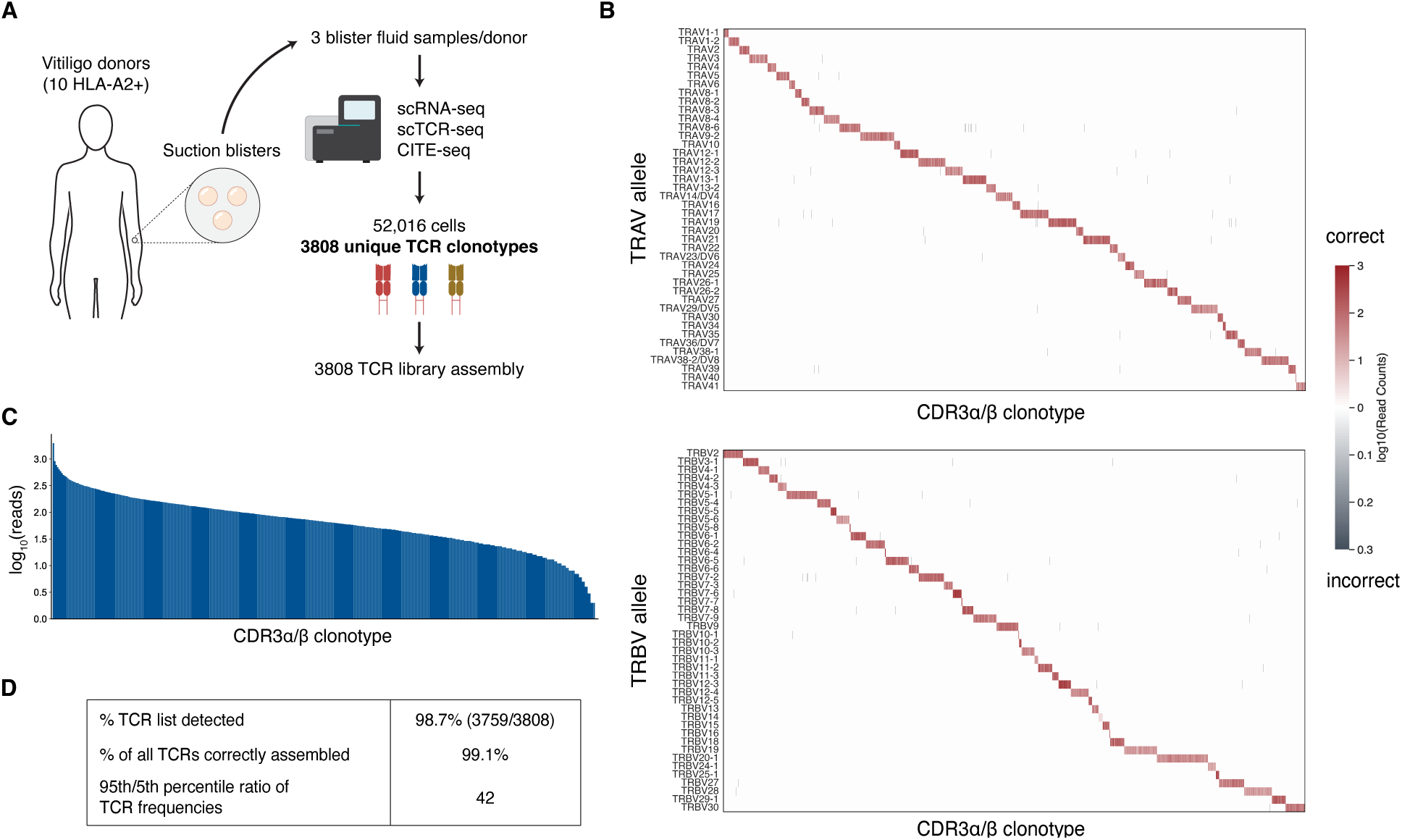
Pooled assembly of 3,808 TCRs from scTCR-seq of vitiligo blister fluid. **(A)** Sample origin and extraction of 3,808 TCRαβ clonotypes. Blister fluid samples from 10 HLA-A2^+^ vitiligo donors were analyzed via 10X scRNA-seq, scTCR-seq, and CITE-seq to capture 3,808 unique TCR clonotypes with matched gene expression data. (B) Heatmap depicting accuracy of TRAV and TRBV alleles pairing to CDR3α and CDR3β sequences, respectively, as log_10_-transformed read counts of all TRAV-CDR3α and TRBV-CDR3β combinations. Red indicates correct pairing and grey indicates incorrect pairing. **(C)** Log_10_-transformed read counts of complete, correctly assembled TCRs. **(D)** Summary data on the proportion of 3,808 TCRs detected, proportion of all TCRs correctly assembled, and frequency distribution.

We assembled these 3,808 vitiligo-associated TCRs using TCRAFT with greater than 1000-fold coverage at each step. We analyzed the assembled TCR library via Oxford Nanopore long-read sequencing to evaluate correct pairing between TCR components and assess the TCR frequency distribution. Sequencing revealed highly accurate pairing between CDR3α and CDR3β sequences and their corresponding TRAV and TRBV alleles (**Figures 2B, S3, and S4**). 99.1% of sequences in the final product consist of correctly paired TRBV, TRAV, CDR3α, and CDR3β sequences corresponding to the 3,808 TCR list obtained from scTCR-seq (**Figure 2B**). We detected 3,749 of 3,808 TCRs (98.7%) via long-read sequencing with a ∼42-fold variation in TCR frequency between the 5^th^ and 95^th^ percentiles (**Figures 2C and 2D**). We complemented this analysis with short-read sequencing of both the TRAV-CDR3α and TRBV-CDR3β regions at greater sequencing depth and detected 99.9% of both CDR3α and CDR3β sequences (3,804/3,808).

### Library-on-library TCR-antigen screen using pMHC-pseudotyped lentiviruses

To screen these TCRs for reactivity, we expressed our 3,808 vitiligo-associated TCR library in a clonal TCR-null Jurkat J76 T cell line encoding an NFAT-CFP reporter^35^. Most approaches to screen TCRs against antigens for reactivity are limited to individual antigens or fail to generate data on interacting TCR-antigen pairs. We and others have demonstrated the feasibility of using pMHC-displaying lentiviruses to infect antigen-specific TCR-expressing cells and extract paired TCR-antigen information with single-cell sequencing via the approaches RAPTR and ENTER-seq^28,36^. While in principle allowing for high-throughput antigen screening, our original RAPTR platform relied upon pMHC tetramers to pre-enrich TCRs from large libraries. Noting that pMHC-pseudotyped lentiviruses potently activate TCR-expressing cells, we optimized RAPTR to extract paired TCR-antigen data for 3,808 TCRs against 101 HLA-A2-binding antigens in one step using a highly sensitive NFAT reporter^35^ (**Figure 3A**). By capturing cells activated by pMHC viruses, we identify antigen-reactive cells based on function rather than solely viral infection, significantly reducing noise from non-specific transduction. This high selectivity is essential for screening large TCR libraries and eliminates laborious pre-enrichment steps.

**Figure 3.**
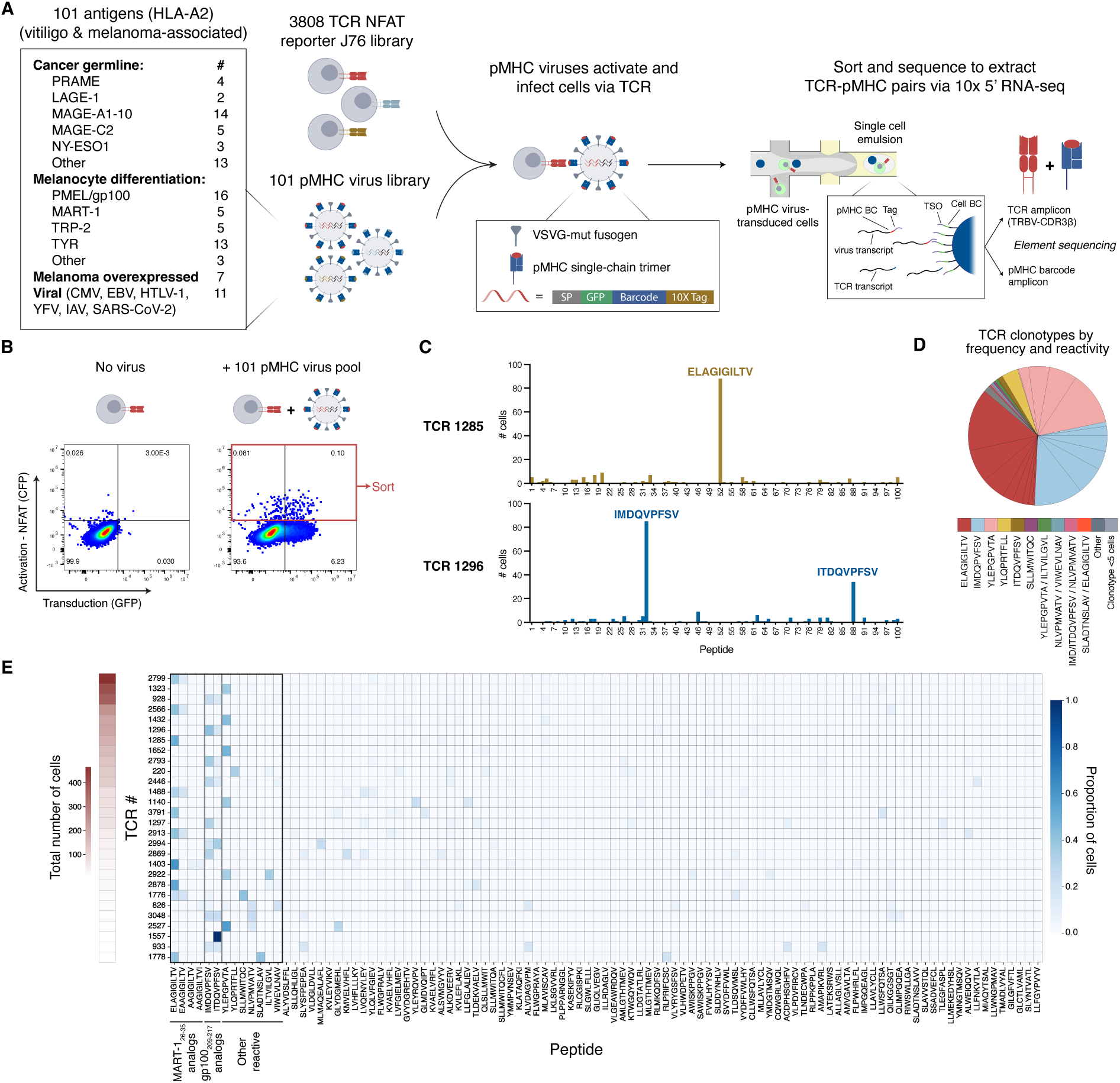
RAPTR enriches and pairs reactive TCRs with antigens. **(A)** Schematic of TCR library screen with 101 pMHC-pseudotyped lentiviruses to extract TCR-antigen pairs. 3,808 TCR-expressing NFAT reporter J76 cells are mixed with a 101-pMHC virus library containing vitiligo, melanoma, and viral antigens. Cells activated and transduced by the virus library are sorted and single-cell sequenced to capture reactive TCRs and cognate antigens. **(B)** Cells activated (CFP^+^) and transduced (GFP^+^) by 101-pMHC virus library. Activated CFP^+^ cells and CFP^+^GFP^+^ cells are sorted to capture both TCRs reactive to pMHC virus and virally integrated pMHC barcodes. CFP^+^GFP^+^ cells are sequenced via 10X single-cell sequencing. **(C)** Bar plots for two enriched TCRs (TCR #1285 and #1296 as examples) showing the number of cells assigned each pMHC identity from single-cell sequencing. Each cell is assigned the pMHC identity with the highest UMI count. Each TCR clonotype is annotated as reactive to pMHCs with the most associated cells. TCR #1285 is reactive to ELAGIGILTV (MART-1) while TCR #1296 is reactive to IMDQVPFSV (gp100) and ITDQVPFSV (gp100). **(D)** Summary of reactive TCR clonotypes showing frequency (wedge size) and antigen reactivity (color) of each clonotype. **(E)** Heatmap depicting the distribution of pMHC assignments for each reactive TCR clonotype, sorted by total TCR frequency (*y* axis). Each row represents a TCR clonotype and values correspond to the sample data depicted in (C), scaled to the total number of cells. MART-1_26-35_ and gp100_209-217_ analogs, as well as other reactive epitopes are grouped on the left side of the heatmap.

We constructed a list of 101 HLA-A2-binding epitopes composed of melanoma-associated antigens^37^, vitiligo-associated antigens in the Immune Epitope Database (IEDB)^38^, and known viral antigens from CMV, EBV, HTLV-1, YFV, IAV, and SARS-CoV-2 (**Table S3**). We then assembled a RAPTR library, with each epitope represented by a pMHC-pseudotyped lentivirus containing a matched barcode to enable integration into bulk and single-cell sequencing workflows (**Figure S5**). Improving upon the prior RAPTR workflow, we used the 101-pMHC virus library to both activate and transduce antigen-specific cells. After stimulating the 3,808 TCR library with the 101-pMHC virus library, we sorted activated TCR-expressing cells expressing an NFAT reporter (CFP^+^), a subset of which were transduced (GFP^+^) (**Figure 3B**).

As a feature of our assembly method, we codon-optimized CDR3α/β oligos to be fully unique, allowing complete TCR identities to be determined by sequencing either CDR3α or CDR3β. To extract paired TCR-pMHC information, we performed single-cell RNA sequencing on the transduced cell subset using 10X Genomics GEM-X 5’ chemistry. We included a targeted primer at the reverse transcription step and developed a custom amplification protocol to capture the TCRβ chain and identify complete TCR clonotypes. We additionally included a 10X Genomics capture tag sequence in the viral genome analogous to ENTER-seq^36^. This capture tag enables viral genomes encoding pMHC barcodes to be directly captured by the template switch oligo (TSO) sequence on 10X 5’ gel bead-in emulsion (GEM) beads.

After filtering for doublets, we recovered 3,202 cells with paired TCR and pMHC information. Each cell was assigned a TCR and pMHC identity based on the ratio of unique molecular identifiers (UMIs) for the dominant TCR or pMHC relative to all TCR or pMHC UMIs for that cell. Each TCR identity was then be paired to a cognate pMHC by examining the pMHC assignments of all corresponding cells expressing a given TCR. For example, TCR #1285 recognizes a MART-1_26-35_ analog, ELAGIGILTV, while TCR #1296 recognizes two gp100_209-217_ analogs, IMDQVPFSV and ITDQVPFSV (**Figure 3C**). Collectively, a total of 28 candidate TCR clonotypes were captured with at least 5 total cells per clonotype, predominantly annotated as recognizing MART-1_26-35_ analogs, gp100_209-217_ analogs, and gp100_280-288_ (**Figures 3D and 3E, Table S4**). We observed a strong association between cells infected by viruses displaying MART-1_26-35_ analogs ELAGIGILTV and EAAGIGILTV, as well as gp100_209-217_ analogs ITDQVPFSV and IMDQVPFSV. Recovering a high quantity of cells increases confidence in assigning a pMHC identity to a given clonotype but TCR-antigen pairs can be unambiguously established with as few as 6 cells. Some low frequency (< 1%) clonotypes were assigned 2-3 possible pMHC identities, indicating a potential limit of detection.

### Detection and validation of antigen-reactive TCRs with peptide-pulsed APCs

We next screened this 3,808 vitiligo TCR J76 library against peptide-pulsed antigen-presenting cells to validate putative antigen-reactive TCR clonotypes captured by our single-cell RAPTR workflow, demonstrate the compatibility of our TCR assembly strategy with a well-precedented and widely used antigen discovery method, and capture antigen-reactive TCRs against a broader array of pMHCs (**Figure 4A**). We generated a list of 561 antigens consisting of known vitiligo epitopes from the IEDB (36), an expanded list of melanoma-associated antigens (195)^37,39,40^, differentially expressed genes in diseased melanocytes filtered for binding to HLA-A2 using NetMHCpan4.1^41^ (226), and viral antigens (104) (**Table S3**). The 101 antigens represented by the RAPTR pMHC virus library are a subset of this broader 561-antigen list.

**Figure 4.**
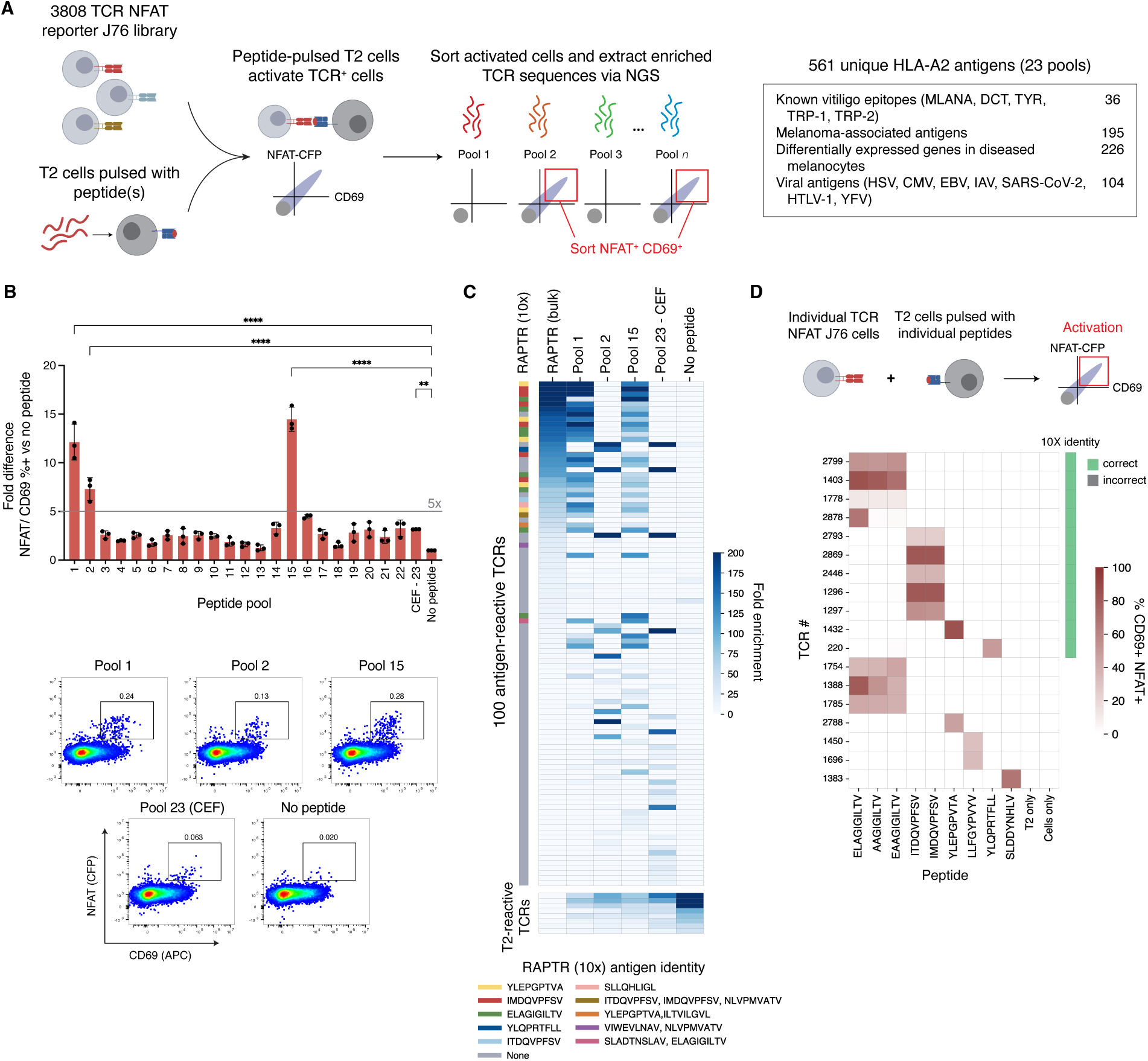
Expanded reactivity screen of 3,808 TCRs and validation with antigen presentation assay. **(A)** Schematic of screening approach. 3,808 TCR-expressing NFAT reporter J76 cells are cocultured with HLA-A2^+^ T2 cells pulsed with 23 pools of peptides (561 unique antigens). Activated (NFAT^+^CFP^+^) cells are sorted and sequenced via next-generation sequencing (NGS) to capture reactive TCRs. **(B)** Bar plots and representative flow plots of cells activated by peptide pools. Bars report the ratio of % cells NFAT^+^CD69^+^ for peptide pools versus the no peptide control. Grey line indicates 5× threshold for sorting cells activated by non-CEF peptide pools. Data shown as mean ± S.D. for 3 replicates. **(C)** Heatmap depicting TCRs >10-fold enriched by 101 virus pool (RAPTR) or indicated peptide pool. Fold enrichment was calculated by dividing the frequency of each TCR in sorted cells with starting frequency in the naïve 3,808 TCR library. Several TCRs enriched in both no peptide (T2 cells only) and peptide pool conditions, shown in the second heatmap; these TCRs are not considered to be antigen-reactive. RAPTR (bulk) represents NGS analysis of cells activated but not necessarily transduced by 101-pMHC virus library. RAPTR (10X) bar indicates classification of antigen reactivity for TCRs identified in single-cell workflow (Figure 3). **(D)** Heatmap showing activation (% NFAT^+^CD69^+^) of individual TCR-expressing NFAT J76 cells with individual peptide-pulsed T2 cells for TCR clonotypes identified via RAPTR or peptide pool screens. *P* values are calculated with a one-way ANOVA (unpaired) with Dunnett’s correction for multiple comparisons. \*\*\*\**P* < 0.0001; \*\*\**P* < 0.001; \*\**P* < 0.01; \**P* < 0.05; n.s., *P* ≥ 0.05.

Upon stimulating the vitiligo TCR J76 library with TAP-deficient HLA-A2^+^ T2 cells pulsed with 23 pools encompassing 561 unique peptides, we observed TCR activation (NFAT^+^ CD69^+^) in response to several pools (**Figures 4B and S6**). Vitiligo- and melanoma-associated peptide pools #1, 2, and 15 exhibited the most significant activation of the 23 pools, stimulating greater than five-fold the small proportion of cells activated by the no-peptide negative control. We sorted cells activated (NFAT^+^ CD69^+^) by these peptide pools as well as peptide pool #23, which consisted of 96 common HLA-A2-binding viral peptides. We then PCR-amplified the TCRβ chain from genomic DNA to capture CDR3β sequences of enriched TCRs by next-generation sequencing. Noting that the RAPTR library is a potent antigen-specific activator of T cells, we additionally set out to determine the concordance between the TCRs captured by the single-cell RAPTR workflow and those activated by the viral library. To do so, we sorted and bulk sequenced all T cells activated by the 101 RAPTR pMHC virus library.

We compared TCR frequencies in enriched cells against the naïve 3,808 TCR J76 cell line to identify antigen-reactive TCRs. Across all stimuli, 100 unique TCR clonotypes enriched more than 10-fold compared to the naïve library after filtering TCRs enriched in the no peptide control (**Figure 4C, Table S5**). 47 TCR clonotypes were unambiguously reactive to melanocyte antigens while 25 clonotypes reacted to viral antigens. Peptide pools #1 and #15, as well as RAPTR, contained overlapping peptides including MART-1_26-35_ analogs, gp100_209-217_ analogs, and gp100_280-288_. The strong concordance between TCR clonotypes enriched by these peptide pools and RAPTR affirms our ability to extract reactive TCR clonotypes with orthogonal antigen discovery methods. The reactive TCRs include 26 of 28 TCR clonotypes detected in our single-cell RAPTR workflow. The two absent clonotypes were low frequency in the single-cell analysis, representing only 0.37% of cells. We additionally noted several TCR clonotypes enriched by all peptide pools and the no peptide control. These TCRs are reactive to T2 cells but not the RAPTR pMHC virus library, highlighting a potential strength of pMHC-pseudotyped lentiviruses. Using these orthogonal antigen discovery approaches, we demonstrate that we can enrich rare antigen-specific TCRs from libraries of thousands of TCRs using both virally displayed pMHCs and co-culture with peptide-pulsed antigen-presenting cells.

To validate hits, we established individual clonal TCR J76 cell lines for a selection of TCRs and verified reactivity to individually peptide-pulsed T2 cells (**Figure 4E**). Of these, all TCR-antigen pairs identified via single-cell sequencing align with individual T2-peptide stimulation data. By integrating large-scale TCR assembly and reactivity screening with pMHC-displaying viruses and peptide-pulsed APCs, we enable large-scale screening of TCRs from tissue-derived TCRs.

### Phenotypes and antigen reactivity of blister fluid TCRs in vitiligo

We next sought to examine the gene expression programs of antigen-specific TCRs in vitiligo using our single-cell RNA-sequencing (scRNA-seq) dataset. Antigen-specific T cells have been reported to exhibit tissue-resident memory (T_RM_)^42^ and cytotoxic phenotypes^32^, yet the functional interplay and contributions of these cell states to vitiligo pathogenesis remain unclear. Paradoxically, the same T cell phenotypes implicated in autoimmunity are broadly protective in melanoma and other cancers^43,44^. Although vitiligo and melanoma share antigenic targets^7,14–17^, it remains unclear whether antigen-specific T cells adopt similar transcriptional programs in both contexts.

To address these questions, we created an atlas of T cell phenotypes for all 10 donors via unsupervised clustering (**Figures 5A and S9**). Most cells captured were CD8^+^; a minority of cells were regulatory T cells, as evidenced by expression of *CD4* and *FOXP3* (**Figures 5B and S9**). Among CD8^+^ T cells, we observed a ‘Cytotoxic CD8’ cluster defined by high expression of multiple granzymes and perforin (*PRF1*), consistent with prior literature highlighting the role of cytotoxic CD8^+^ T cells in vitiligo^45^. This phenotype was corroborated by high expression of HLA-DR in this cluster (**Figure 5B**). We further detected two clusters characterized by expression of *ZNF683 (HOBIT)*, a key transcription factor marking tissue-resident CD8^+^ T cells, and the chemoattractant *GPR183 (EBI2)*, marking CD8^+^ T cells in an activated and migratory state^46^.

**Figure 5.**
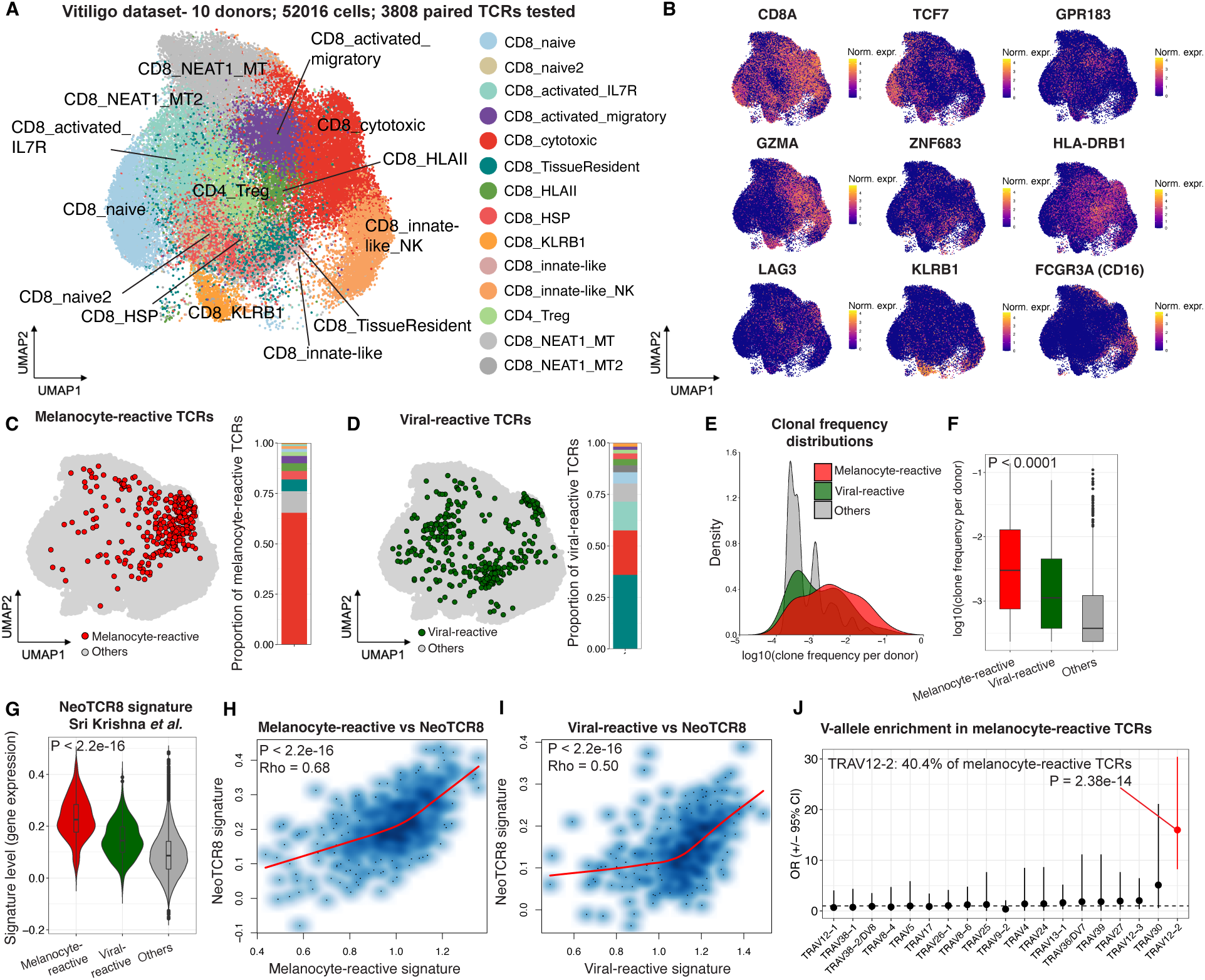
Phenotypes of TCRs with detected antigen reactivity. **(A)** UMAP representation of T cell clusters from integrated single-cell RNA-seq (scRNA-seq) analysis of 10 vitiligo donors. **(B)** Expression of key markers corresponding to clusters in (A). **(C-D)** Cluster distribution of antigen-detected TCRs for melanocyte-reactive TCRs (C) and viral-reactive TCRs (D). **(E-F)** Clonal frequency distribution of antigen-detected TCR clonotypes vs non-antigen-detected TCR clonotypes (‘Others’). Clone frequency calculated per donor. Data show antigen-detected clonotypes have a higher clonal frequency compared with non-antigen-detected clonotypes. **(G)** Application of NeoTCR8 signature from Sri Krishna et al.^75^ to clonotypes in our dataset. **(H)** Correlation of melanocyte-reactive gene expression signature with the NeoTCR8 signature across all cells with antigen-reactive TCRs. **(I)** Correlation of the viral-reactive TCR gene expression signature with the NeoTCR8 signature across all cells in (A). **(J)** V-allele enrichment in melanocyte-reactive TCRs compared with non-antigen-detected TCRs showing enrichment of the TRAV12-2 allele. Red indicates adjusted *P* < 0.05 by Fisher’s exact test.

To resolve the identities of these clusters and compare them to known T cell phenotypes in vitiligo, we surveyed expression of effector molecules, cytokines, chemokines, and transcription factors across the entire T cell atlas. The ‘Cytotoxic CD8’ T cell cluster expressed high levels of multiple granzymes, including *GZMA*, *GZMB*, *GZMH*, and *GZMM*, but did not display high expression of *GZMK* (**Figures 5B and S9**), in contrast to the CD8^+^ GZMK^+^ cells previously shown to drive pathogenesis in rheumatoid arthritis^47^ and airway inflammation^48^. Suggesting an exhausted state, the ‘Cytotoxic CD8’ cluster expressed *PDCD1*, *CTLA4*, and *TIGIT* (**Figure S9**). Among cytokines and chemokines, *IFNG* and *CCL4*/5 were predominantly expressed, in addition to *IL32* (**Figures 5B and S9**), which previous studies link to effector T cell responses in skin inflammation^49–51^. The tissue-resident cluster was broadly *HOBIT^+^ CD69^+^ ITGAE (CD103)^+^ CD49A (ITGA1)^-^,* consistent with prior reports^52–54^. Although these genes were most highly expressed in our ‘Tissue Resident’ cluster (**Figure S9, Table S6**), both *CD69* and *ZNF683* were also detected at a lower level across the T cell atlas (**Figures 5B and S9**). This aligns with observations that CD8^+^ T cells generally adopt a tissue-resident memory phenotype in vitiligo^52–54^.

We next paired TCR-seq with the scRNA-seq data (**Table S6**) to characterize the phenotypes of clonally expanded and antigen-reactive TCRs. A total of 309 melanocyte-reactive T cells, representing 47 TCR clonotypes (**Table S7**), were distributed over multiple clusters in the T cell atlas, with the majority localized to the CD8^+^ cytotoxic cluster (**Figure 5C**). In contrast, the 329 viral-reactive T cells, representing 25 distinct TCR clonotypes (**Table S7**), were predominantly located in the CD8^+^ tissue-resident cluster, with comparatively less enrichment in the cytotoxic cluster (**Figure 5D**). Melanocyte-reactive TCRs exhibited higher clonal frequencies across and within individual donors compared to viral-reactive TCRs and non-antigen-detected TCRs (‘others’) (**Figures 5E, 5F, and S10**). These findings underscore the capacity of our approach to detect TCRs specific to disease-associated antigens.

The presence of effector and exhaustion markers on antigen-specific T cells in vitiligo prompted us to assess their transcriptional similarity to T cells in melanoma. Given that we identified TCRs reactive to shared vitiligo and melanoma antigens, we applied a previously published gene expression signature of antigen-specific CD8^+^ T cells (NeoTCR8) in melanoma^55^ to our dataset. This signature was markedly elevated in antigen-reactive cells, especially for melanocyte-reactive TCRs (melanocyte vs. other, P < 2.2e-16) (**Figure 5G**). We then derived transcriptional signatures of melanocyte- and viral-reactive TCRs using differential gene expression analysis^55^ (**Tables S13 and S14**), both of which positively correlated with NeoTCR8 (**Figures 5H and 5I**). Applying additional signatures of melanoma-reactive TCRs from Oliveira et al.^7^ to our dataset, we noted the strongest enrichment of terminally and progenitor exhausted (TTE and TPE) signatures in melanocyte-reactive T cells from our dataset (**Figure S11A**). We further observed correlation of our melanocyte- and viral-reactive signatures with all Oliveira et al.^7^ melanoma signatures **(Figures S11B and S11C)**. Collectively, these data suggest shared CD8^+^ T cell transcriptional phenotypes across autoimmunity, cancer, and infection. The melanocyte- and melanoma-reactive signatures shared genes for exhaustion markers (*PDCD1, LAG3, TOX, ENTPD1*). *CD70* and *CD27*, both implicated in cancer immune evasion^56^, were also shared by melanocyte and melanoma signatures but are notably absent from the viral-reactive signature (**Tables S13 and S14**).

Previous studies have reported biased germline V allele usage in antigen-specific TCRs in autoimmunity^57^. To investigate whether antigen-specific TCRs are similarly restricted in vitiligo, we compared individual V or J allele usage in melanocyte-reactive TCRs with detected antigen relative to other TCRs. TRAV12-2 emerged as the only significantly enriched allele (**Figure 5J**), likely reflecting reactivity to MART-1 antigens, as previously noted in HLA-A2^+^ melanoma patients^58^. To probe potential shared TCRs reactive against MART-1 in vitiligo and melanoma, we compared our TRAV12-2^+^ MART-1-reactive TCRs (**Table S7**) with those catalogued in VDJDb^59^. We found only a single shared chain and no sequence homology to well-established DMF4/5 MART-1-reactive TCRs^60^.

### Assembly and antigen reactivity screening of a 30,000+ TCR library

With a validated methodology to construct TCR libraries, we next set out to determine if we could efficiently assemble tens of thousands of TCRs in a single reaction workflow. We compiled 30,810 TCR sequences derived from tumor-infiltrating lymphocytes (TILs) from surgically resected tumors and matched PBMCs from 21 donors with pancreatic ductal adenocarcinoma (PDAC), as well as PBMCs from 9 PDAC and 2 healthy donors^61,62^ (**Figure 6A, Table S10**). We ordered CDR3α/β oligos at a cost of $0.30 per TCR and assembled the TCR library using TCRAFT. Analogous to the 3,808 TCR library, we characterized the product with both Oxford Nanopore long-read sequencing and short-read Element sequencing. CDR3α and CDR3β sequences paired with corresponding TRAV and TRBV alleles with exceptional accuracy: 99.1% of assembled sequences correspond to a correctly assembled TCR from the library and 98.6% (30,681/30,810) of TCRs were detected via long-read sequencing (**Figure 6B**). TCR frequencies spanned ∼36-fold between the 5^th^ and 95^th^ percentile, a modest distribution suitable for a variety of screening methods (**Figures 6C and 6D**). Short-read sequencing captured 99.9% of CDR3α and CDR3β sequences (30,772/30,810).

**Figure 6.**
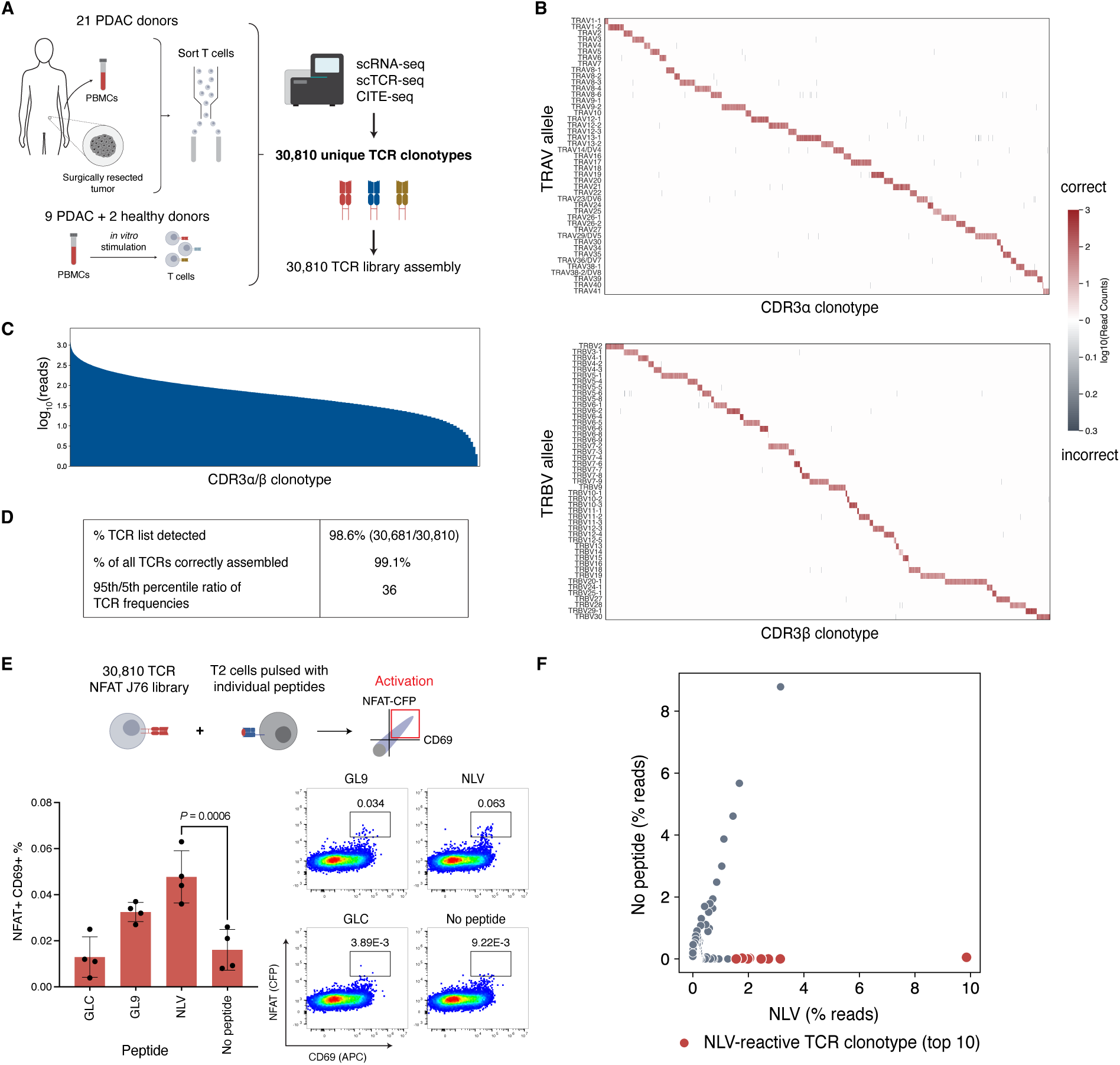
Pooled assembly of 30,810 TCRs from scTCR-seq of PDAC tumor resections and PBMCs. **(A)** Sample origin and extraction of 30,810 TCRαβ clonotypes. **(B)** Heatmap depicting accuracy of TRAV and TRBV alleles pairing to CDR3α and CDR3β sequences as log_10_-transformed read counts of all TRAV-CDR3α and TRBV-CDR3β combinations. Red and grey indicate correct and incorrect pairing, respectively. **(C)** Log_10_-transformed read counts of complete, correctly assembled TCRs. **(D)** Summary data on the proportion of 30,810 TCRs detected, proportion of all TCRs correctly assembled, and frequency distribution. **(E)** Screen of 30,810 TCR library with individually peptide-pulsed T2 cells. Bar plot shows activation as percentage of cells NFAT^+^CD69^+^ with representative flow plots for each antigen. Data is represented as mean ± S.D. for *n* = 4 replicates. *P* values were calculated with a one-way ANOVA (unpaired) with Dunnett’s correction for multiple comparisons. **(F)** Scatterplot representing TCR clonotypes enriched by T2 cells pulsed with NLV or no peptide (negative control). Enriched cells were processed via NGS to extract TCR clonotypes. TCR frequencies are depicted with each dot representing a unique TCR clonotype. Red dots indicate the top 10 TCR clonotypes enriched exclusively by NLV peptide.

To evaluate our ability to isolate rare antigen-reactive TCRs from this large TCR library, we expressed the 30,810 TCR library in a clonal NFAT-CFP reporter J76 T cell line. To experimentally validate that the library was functional, we activated the TCR library with T2 cells pulsed with individual seroprevalent HLA-A2-binding epitopes: influenza-derived GL9 (GILGFVFTL), CMV-derived NLV (NLVPMVATV), and EBV-derived GLC (GLCTLVAML) and sorted activated cells for TCR sequencing analysis. Five donors with available haplotypes were HLA-A*02:01^+^ and, considering the mixed TIL-PBMC library composition, we anticipated a small proportion of the TCRs to be potentially reactive to these immunodominant epitopes. We captured reactivity to GL9 (P = 0.0505) and NLV (P = 0.0006) by a small fraction of cells (**Figure 6E**). We then sorted cells reactive to both T2 cells pulsed with NLV peptide and T2 cells without peptide as a negative control, and sequenced CDR3β amplicons to identify activated TCRs. Of the ten most abundant clonotypes enriched by NLV but not the no-peptide control (**Figure 6F**), six contained α or β chains documented as NLV-specific by data in VDJDb^59^. Two of these clonotypes match TCRs known to recognize NLV (**Table S11**) and three clonotypes are from a donor confirmed to be HLA-A*02:01^+^^63,64^. Our approach to assembling large synthetic TCR libraries enables the sensitive identification of low-frequency antigen-reactive TCRs.

## Discussion

Scalable, cost-effective synthesis and functional screening of TCR libraries with high accuracy is essential to decoding TCR-antigen specificity. We have integrated single-cell CD8^+^ T cell profiling, pooled low-cost assembly of thousands of TCRs, and library-on-library TCR-antigen screening to build a broadly adoptable TCR-antigen screening pipeline and to examine transcriptomic signatures of antigen-reactive TCRs. Our TCR assembly approach, TCRAFT, enables low-cost, pooled synthesis of tens of thousands of TCRs with > 99% assembly accuracy while maintaining native α/β pairing. Immortalized TCR-expressing cells facilitate deep profiling of irreplaceable samples, enable rounds of enrichment, and allow for TCR-antigen pairing in a single step using RAPTR. By integrating TCRAFT with an optimized RAPTR pipeline, we achieve efficient, one-pot screening of TCR-antigen interactions, allowing identification of antigen-specific TCRs from thousands of TCRs with matched gene expression data. Using this pipeline, we screened 3,808 TCRs derived from vitiligo lesions of 10 donors against 101 antigens in a library-versus-library format. We further synthesized a 30,810-TCR library in a single reaction and demonstrated the compatibility of both TCR libraries with peptide-pulsed antigen-presenting cell screening. By linking antigen specificity to scRNA-seq data for vitiligo lesion-associated TCRs, we identified signatures of both melanocyte- and viral-reactive TCR and shared transcriptional phenotypes with antigen-specific T cells in melanoma.

Advances in single-cell and computational methods for sequencing paired TCR chains have produced large TCR datasets with little data on antigen specificity. Fewer than 1 million unique TCR-antigen pairs are known, with less than 4% of these TCRs include both TCR α and β chains^2^. Current pipelines for reconstructing and evaluating TCR sequences for specificity are largely confined to research groups with specialized expertise and typically focus on individual TCRs selected based on clonotypic expansion or phenotype. Although large-scale TCR synthesis and screening would provide unbiased antigen reactivity profiling, existing strategies are expensive, scale-limited, or difficult to execute. Our approach, TCRAFT, is affordable (<$1/TCR), freely available, easy to implement, and compatible with an array of antigen discovery methods. We anticipate that these features will enable broad use, leading to a significant increase in the number and diversity of known TCR-antigen pairs. This will both accelerate the identification of clinically relevant TCRs for adoptive cell therapy and engineered therapeutics and provide valuable training data for computational models of TCR specificity, enhancing predictive frameworks for antigen recognition.

Characterizing antigen-specific T cells in autoimmunity is crucial to decoding disease pathogenesis, developing diagnostic markers, and designing targeted immunotherapies. We therefore applied this pipeline to screen TCRs isolated from vitiligo suction blister fluid, a non-renewable tissue sample. By capturing TCR and gene expression via single-cell sequencing, we screened 3,808 TCRs against a total of 561 antigens and linked TCR reactivity to phenotype.

Analysis of paired scRNA-seq and TCR-seq data from vitiligo blister fluid revealed that melanocyte- and viral-reactive TCRs span multiple transcriptional states. Underscoring the specificity and sensitivity of our screening pipeline, TCRs identified as vitiligo antigen-specific exhibited pronounced clonal expansion. These T cells primarily adopted a tissue-resident phenotype marked by high expression of *CD69, HOBIT,* and *CD103*, or a cytotoxic state characterized by expression of granzymes, *IFNG, CCL3/4/5, PRF1*, and exhaustion markers such as *PDCD1*. Gene signature analysis of antigen-reactive T cells in our dataset revealed significant correlation with antigen-specific T cells in melanoma. The molecular mechanisms driving convergent phenotypes of T cell cytotoxicity and exhaustion in autoimmunity and cancer remain to be determined^65,66^. Our data suggest that T cell exhaustion as a common phenotype of antigen-specific T cells across diseases. Notably, genetic analyses have associated vitiligo with reduced incidence of melanoma^67^, raising the possibility that antigen-reactive T cells in one setting may confer protection in another.

Paired with new methods to capture TCR sequences such as TIRTL-seq^19^, our approach for large-scale TCR assembly and screening completes a pipeline to generate TCR specificity data at an unprecedented scale. Given extensive work expressing TCRs in both primary and Jurkat cells, we expect that our TCR library assembly approach will be compatible with any workflow utilizing TCR-expressing cells^27,35,68–71^. By enabling rapid and cost-effective TCR assembly and seamlessly integrating TCR libraries with multiple antigen screening methods, this pipeline has the potential to advance immunotherapy, accelerate vaccine design, and deepen our understanding of TCR recognition.

## Methods

### Media and cells

HEK293T cells (ATCC CRL-11268) were cultured in DMEM (ATCC) supplemented with 10% fetal bovine serum (FBS; Atlanta Biologics) and penicillin-streptomycin (Gibco). Jurkat J76 T cells, a Jurkat E6.1 cell line devoid of TCR alpha and beta chains, were a gift from M. Heemskerk^35^. J76 cells and T2 cells (ATCC CRL-1992) were cultured in RPMI-1640 (ATCC) supplemented with 10% FBS and penicillin-streptomycin.

### Plasmid cloning and construction

Primer oligonucleotides were ordered from IDT (listed in **Table S12**), gene fragments were synthesized by Twist Biosciences and IDT, oligonucleotide pools were synthesized by Twist Biosciences (250-300 nt) and IDT (300-350 nt), and TCRAFT vectors were constructed by Genscript and Twist Biosciences. psPAX2.1 and pMD2.G were gifts from D. Trono (Addgene #12260 and #12259). pMD2.G-VSV-G-mut is available on Addgene as plasmid #182229^28^. TRBV-TRAC and TRBC-TRAV TCRAFT vectors were designed using native human sequences for TRBV and TRAV alleles from IMGT^72^. Vectors were assembled by Twist Biosciences: TRBV-TRAC in a pGGA cloning vector with chloramphenicol resistance and TRAC-P2A-TRAV in the pTwist Kan HC v2 cloning vector with kanamycin resistance (**Table S1**). A lentiviral expression vector (pHIV backbone) was constructed by inserting a gene fragment (IDT) encoding LacZ flanked by SapI recognition sites (5’-GCA-SapI site-LacZ-SapI site-ATC-3’) and removing an undesired backbone SapI site with the Quikchange Lightning Site-Directed Mutagenesis Kit (Agilent #210518).

To generate pMHC-displaying lentiviruses, 101 pMHCs were cloned into a pHIV backbone by Genscript as single-chain trimers (SCT)^28^ (HGH signal peptide-peptide-G4S linker-β2 microglobulin-HLA-A*02:01) with two cysteine mutations to stabilize peptide-MHC binding (Y84C in HLA-A2 and G2C in the G4S linker). Our previously described pLeAPS-GFP plasmid^28^ was modified to include a 10X bead-compatible tag sequence (TSO)^36^ and an extended 18 bp barcode via randomized primers.

Individual TCRs for validation were ordered as TRBV-TRBC-P2A-TRAV gene fragments (IDT) and assembled via Golden Gate into the same pHIV backbone described above, modified to contain the TRAC flanked by BsmBI sites.

### Selection of TCRAFT Golden Gate overhang sets

Unique, highly orthogonal overhangs were designed for each TRBV-TRAC (n = 48) and TRBC-TRAV (n = 45) vector to ensure each CDR3α and CDR3β pairs with correct alleles in hierarchical Golden Gate assembly reactions while preserving coding sequences. To design optimal overhangs, a dataset of ligation fidelities for all 256 possible 4-bp overhangs was leveraged^29^. All possible 4-bp overhangs were identified in the last 8 amino acids of each native TRBV and TRAV sequence and the first 10 amino acids of the TRAC and TRBC using combinatorial codon shuffling. 4-bp overhangs for TRBV-TRAC and TRBC-TRAV vectors were selected using two custom scoring metrics for assembly efficiency: one to reduce off-target pairing and one maximizing correct circular product assembly. An overhang set 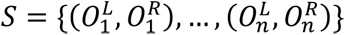 was defined as a set of *n* overhang pairs 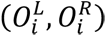, where 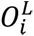 and 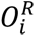 correspond to the left and right 4-bp overhangs assigned to vector *i,* respectively. Using the ligation fidelity dataset^29^, the probability of correct overhang pairing was computed for any 4-bp overhang *o*_*i*_ using the formula *p*(*o*_*i*_) = *N*_*correct*_/*N*_*total*_, where *N*_*correct*_ is the number of reads in which *o*_*i*_ ligates correctly to its complementary Watson-Crick pair, and *N*_*total*_ is the total number of reads associated with overhang *o*_*i*_ in the dataset.

The first overhang set scoring function *f*_1_(*S*), which optimizes for maximal overhang orthogonality, was computed by taking the product of the correct assembly probability across all overhangs in the set as follows:

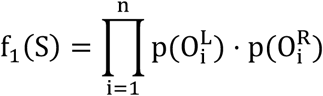

The second overhang set scoring function *f*_2_(*S*), which optimizes for maximal correct circular assembly, was computed by taking the minimum of the correct assembly probability for each individual vector. Aggregating scores across each vector using the minimum instead of calculating an average or sum ensures that correct circular assembly values across vectors all exceed a minimum threshold:

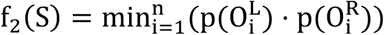

The final scoring function used in the overhang set optimizer was a weighted average of *f*_1_(*S*) and *f*_2_(*S*) to balance both scoring objectives. A Markov Chain Monte Carlo (MCMC) sampling approach was used to stochastically design overhang sets that achieve high scores on the combined overhang set scoring metric. Each round, the optimization algorithms randomly shuffled one of the overhangs in the set, computed the score of the new modified overhang set, and either accepted or rejected this change according to the probability distribution below. *T* = 10^-3^ was empirically selected and the overhang set scoring function converged over 10,000 iterations.

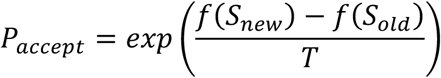

The result was a pair of 4-bp overhangs for each TRBC-TRAV and TRBV-TRAC fragment that preserves coding sequences. A total of four overhang sets were generated: two overhang sets for TRBV-TRAC vectors, and another two sets for TRBC-TRAV vectors, termed A and B for both. TRAV pool A includes TRAV1-1 to TRAV16 (22 alleles) and pool B includes TRAV17 to TRAV41 (23 alleles). TRBV pool A includes TRBV2 to TRBV7-8 (24 alleles) and pool B includes TRBV7-9 to TRBV30 (24 alleles). TRAV and TRBV alleles were split into two pools each to maximize assembly efficiency. To enable broad use, we created a script (below) that accepts TRAV, TRBV, TRAJ, TRBJ, CDR3α, and CDR3β as inputs and generates lists of oligos to order as outputs, split into pools A and B. For convenience, output oligos are split by length (≤ 300 bp and >300 bp) to simplify ordering.

### Validation of Golden Gate overhang sets

Computationally generated overhang sets were experimentally validated. Four pools of backbone library vectors were generated by inserting pools of oligos containing the overhang sets, BsmBI recognition sites, and a defined barcode for each overhang to enable computation of assembly fidelity into pGGAselect (NEB). Oligo pools representing eventual CDR3α/β oligos were flanked with BsmBI recognition sites, prescribed overhangs, and a unique barcode. Oligo pools were amplified according to manufacturer instructions (15 cycles).

Vectors and oligos for each overhang set were pooled in Golden Gate assembly reactions at a 4:1 molar insert-to-vector ratio with 2 μL of BsmBI Golden Gate Enzyme Mix (NEB) and T4 Ligase Buffer (NEB). Golden Gate reactions were cycled (30 cycles of 42C for 5 min, 16C for 5 min; 60C for 5 min, 80C for 20 min). 2 μL of the product was heat-shock transformed into competent Stbl3 *E. coli*, cultured under chloramphenicol (Sigma Aldrich) selection, and midiprepped. Amplicons of ligated insert and flanks were PCR-amplified (30 cycles) and submitted to Genewiz (Amplicon-EZ NGS). Comparison of barcodes adjacent to insert overhangs allowed for quantification of on-target versus off-target vector-insert pairing within each overhang set.

### Synthesis and cloning of TCR libraries

TCR libraries were generated in hierarchical Golden Gate assembly reactions as follows. Oligo pools A and B (Twist for 250-300 nt, 97% of TCRs; IDT for oligos > 300 nt, 3% of TCRs) were generated by the oligo generation script. Oligo pools were re-suspended as directed by the manufacturer. If ordered with separate manufacturers (Twist for ≤ 300 bp and IDT for >300 bp), oligos pools were mixed proportionally prior to amplifying. 20 ng of each oligo pool was PCR-amplified using NEBNext Ultra II Q5 Master Mix (NEB) (98C for 30s; 12 cycles of 98C for 30s, 68C for 30s, 72C for 30s; 72C for 5 min) using Oligo_f and Oligo_r primers followed by a 1.8× SPRIselect bead (Beckman Coulter) cleanup and elution in elution buffer (Thermo Fisher).

In step 1, CDR3α/β oligo pools A and B were inserted into the TRBV-TRAC-containing vectors (A and B). In parallel, TRBV-TRAC vector pool A was reacted with CDR3α/β oligo pool A, and TRBV-TRAC vector pool B was reacted with CDR3α/β oligo pool B. The following were combined to generate two 15 μL reactions (A and B): 0.1 pmol of vector mix, 0.4 pmol CDR3α/β oligo pool, 5 μL NEBridge Ligase Master Mix (NEB), 1 μL BbsI-HF enzyme (NEB), and nuclease-free water (Thermo Fisher). Both Golden Gate reactions were completed: 60 cycles of 37C for 5 min, 16C for 5 min; 65C for 20 min; hold at 4C. Post-reaction, a clean-up cut was performed to ensure complete elimination of unreacted vector by adding 1 μL of BbsI-HF enzyme (NEB) and 1 μL of shrimp alkaline phosphatase (rSAP) (NEB) to the reaction mix and incubating at 37C for 1 hour followed by 65C for 20 minutes for heat inactivation.

The step 1 products were drop dialyzed for 3 hours on an MCE membrane filter (0.025 pore size) (Millipore Sigma). Dialyzed products were electroporated into ElectroMAX® DH10β electrocompetent *E. coli* (Thermo Fisher, Cat No. 18290015) according to manufacturer instructions. Following overnight culture under chloramphenicol (Sigma Aldrich) selection, electroporation efficiency was evaluated, and products 1A and 1B were midiprepped using a NucleoBond Xtra Midi EF kit (Macherey-Nagel). A minimum of 1000× library coverage was achieved at each electroporation step. Step 1A and 1B products were mixed proportionally according to the number of oligos in each oligo pool, henceforth referred to as step 1 product (minimum 300 ng required).

In step 2, step 1 product was reacted with the TRBC-TRAV-containing vectors (A and B) to insert TRBC-P2A-TRAV fragments between CDR3β and CDR3α. For step 2A, the following were combined to generate a 20 μL reaction: 150 ng of step 1 product and TRBC-TRAV pool A at a 1:2 molar ratio (313 ng), 2 μL of T4 ligase buffer (NEB), 1 μL NEBridge BsmBI-HF-v2 Master Mix (NEB), and nuclease-free water (Thermo Fisher). For step 2B, the following were combined to generate a 20 μL reaction: 150 ng of step 1 product and TRBC-TRAV pool B at a 1:2 molar ratio (313 ng), 2 μL of T4 ligase buffer (NEB), 1 μL NEBridge BsaI-v2 Master Mix (NEB), and nuclease-free water (Thermo Fisher). Step 2A and 2B were completed separately with the following cycling conditions: 60 cycles of 42C (BsmBI, step 2A) or 37C (BsaI, step 2B) for 5 min, 60C for 5 min; 80C for 20 min; hold at 4C. Post-reaction, 1 μL of BsmBI-v2 enzyme (NEB) or 1 μL of BsaI-HF-v2 enzyme (NEB) were added to steps 2A and 2B, respectively, in addition to 1 μL of rSAP (NEB). Both were incubated at 55C (step 2A, BsmBI) or 37C (step 2B, BsaI) for 1 hour followed by 80C for 20 minutes. Step 2 products were dialyzed and electroporated identically as in step 1, described above, followed by culture under chloramphenicol (Sigma Aldrich) selection. Midiprepped step 2 products were mixed proportionally to form the complete TCR library (minimum 244 ng required).

In step 3, the complete synthetic TCR library was transferred to a destination lentiviral transfer vector, described above, to enable expression in cells. The following were combined to generate a 15 μL reaction: 0.05 pmol destination vector, 0.1 pmol of mixed step 2 product, 5 μL of NEBridge Ligase Master Mix (NEB), 1 μL of SapI enzyme (NEB), and nuclease-free water (Thermo Fisher). The following cycling conditions were used: 60 cycles of 37C for 5 min, 16C for 5 min; 60C for 5 min; hold at 4C. Immediately before dialyzing, the reaction was incubated at 60C for 5 min and 65C for 20 min to enable heat inactivation. The step 3 products were dialyzed and electroporated identically to steps 1 and 2, described above, followed by culture under carbenicillin (Sigma Aldrich) selection. The final product is a complete, pooled TCR library in a lentiviral expression vector.

### TCR library and oligo pool sequencing

To examine and quantify assembled TCR sequences, TCR plasmid libraries and oligo pools were sequenced. Products of reactions 1, 2, and 3 contain several unique cut sites including *XbaI.* Products were digested with FastDigest XbaI (Thermo Fisher) in FastDigest Buffer (Thermo Fisher) for one hour at 37C. The digested products were purified via columns by mixing products 1:1 with binding buffer from GeneJET Gel Extraction Kit and following kit instructions. Oxford Nanopore libraries were generated via ligation with kit SQK-LSK114 (ONT) and sequenced on a PromethION R10 flowcell (ONT). For the 3,808 TCR library, the intermediate products (reactions 1 and 2) and final product (reaction 3) were sequenced. For the 30,810 TCR library, only the final product (reaction 3) was sequenced. Long-read sequencing enables minimally biased evaluation of correct pairing, as it avoids PCR steps that depend on specific primer binding sites.

For greater read depth, short-read sequencing was used to sequence TRAV-CDR3α regions (primers: TRAV_CDR3A_f, TRAV_CDR3A_r), TRBV-CDR3β regions (primers: TRBV_CDR3B_f, TRBV_CDR3B_r), and CDR3α-CDR3β oligos (primers: Oligo_Illumina_f and Oligo_Illumina_r). All primers were used at 1 μM. PCRs were completed with NEBNext Ultra II Q5 Master Mix (NEB #M0544). CDR3α-CDR3β oligo pools were PCR-amplified to add partial Illumina adapters with the above primers (50 ng template; 98C for 30 sec; 13 cycles of 98C for 30 sec, 70C for 30 sec, 72C for 30 sec; 72C for 1 min). TRAV-CDR3α and TRBV-CDR3β regions were PCR-amplified with above primers binding conserved regions to add partial Illumina adapters. Template for this reaction consisted of plasmid library (50 ng); reaction conditions: 98C for 30 sec; 15 cycles of 98C for 30 sec, 70C for 30 sec, 72C for 45 sec; 72C for 1 min. A second PCR added indices and complete Illumina adapters to all amplicons (primers: Illumina_Truseq_f and Illumina_Truseq_r; 20-100 pg template; 98C for 30 sec; 15 cycles of 98C for 30 sec, 70C for 30 sec, 72C for 30 sec; 72C for 1 min). Amplicons were pooled in an equimolar manner and sequenced with an Element AVITI, generating paired end 300 bp reads.

#### Long-read sequence processing

Long-read sequencing analysis was completed in Python. Reads were filtered with a size cutoff 8500 bp for the final TCR library (step 3, size ∼8800 bp), 2800 bp for step 1 product, and 3500 bp for step 2 product. To filter out non-TCR sequences, reads were filtered for the presence of a TRAC and TRBC. Next, each read was mapped to a TRAV and TRBV and sequences in the adjacent CDR3 regions were extracted. Each CDR3 region was independently searched for the designed CDR3α and CDR3β sequences; in addition to mapping CDR3α and CDR3β regions to the designed list, new CDR3α and CDR3β sequences were also annotated. Correctly paired, incorrectly paired, and new complete TCR sequences were recorded and counted. Downstream analysis and plots were generated with Matplotlib and Seaborn in Python.

#### Short-read sequence processing

Short read sequencing analysis was completed in Python. For CDR3 oligo pool sequencing data, forward and reverse reads were merged into a single contiguous read using the PEAR software package (v0.9.10). Following read merging, reads were annotated by performing an exact match dictionary search against oligo sequences ordered. For TRAV-CDR3α and TRBV-CDR3β amplicons, forward and reverse reads were processed independently. The forward reads, which mostly captured the variable region, were used to annotate the variable region for each read via exact match lookup. The reverse reads, which mostly captured the CDR3, were used to extract and annotate the CDR3 coding sequence via exact match lookup.

### Single-cell RNA and TCR sequencing

#### 3,808 TCR library

##### Suction blister biopsy and processing

Individuals with rapidly progressing, active vitiligo were recruited under an Institutional Review Board–approved protocol (H-14848) at the University of Massachusetts Chan Medical School. Participants with a diagnosis of vitiligo by clinical exam performed by a dermatologist and treatment-naive for more than 6 months were included for sampling. Clinical blistering sites were examined through suction blister biopsies focused on specific areas showing disease symptoms. Blisters were purposefully formed over confetti or trichrome lesions, incorporating both affected and surrounding skin to capture immune cells. Active lesions not characterized as confetti or trichrome were sampled at the border zone between depigmented and pigmented skin. Ten suction blisters were obtained and pooled for each patient, as previously described^73^. The extracted blister fluid was centrifuged at 350g for 10 minutes at 4C, stained with a pool of Immundex dextramers for 30 minutes at room temperature, and washed 3 times with phosphate-buffered saline (PBS). For single-cell sequencing, whole cell pellet from blisters was used for 10X input due to cell counts being significantly lower than maximum cell input recommendation.

##### scRNA-seq library generation

Once the single cell suspension was obtained, samples were processed using the standard Chromium 5’ V1 (VB203 donor) or V2 (others) + TCR library generation workflow (10X Genomics) and sequenced on a NextSeq500/550 (Illumina) instrument following recommended read configurations. BCL files were then converted to FASTQ files using bcl2fatstq (v2.15.1). Reads were filtered, aligned, and quantified using the 10X Cellranger computational suite (vX.3.1) to generate UMI-collapsed gene by cell count matrices.

##### scRNA-seq preprocessing of vitiligo scRNA-Seq and scTCR-Seq data

The Seurat (version 4.4.0) package was used to exclude low-quality barcodes, resulting in a total of 52,016 barcodes that met the following criteria: log10GenesPerUMI (complexity) exceeding 0.75; number of UMIs between 250 and 60,000; number of genes between 200 and 5,500; ribosomal reads less than 30%; and mitochondrial reads less than 20%. Once these high-quality barcodes were obtained, non-PseudoY genes were removed from the dataset, resulting in a total of 20,688 genes analyzed across 52,016 cells from 10 donors. Subsequently, the Seurat package was employed to integrate the filtered gene-by-cell matrices, which were then analyzed using a typical unsupervised procedure including normalization, scaling, dimensionality reduction, batch correction, cell clustering, and differential gene expression analysis.

TCRs were called using CellRanger and filtered to a final set of 3,808, selected for overlap with scRNA-seq data and the presence of a single alpha and beta chain. Clonotypes were defined by V/J gene usage and CDR3 sequences of both chains. Only TCRs with paired transcriptomic data from quality-controlled cells were retained.

##### Signature analyses with scRNA-seq data

To determine signatures of melanocyte-reactive and viral-reactive cells, differential expression analyses of antigen-reactive cells were performed comparing them against all other cells using MAST^74^ with a covariate (latent_vars) for donor. Genes differentially expressed at FDR P < 0.05 and log fold change > 0 were carried forward for signature analyses. To compare our signatures to signatures from Sri Krishna et al.^75^ and Oliveira et al.^7^, the AddModuleScore() function was used in Seurat. Correlation analyses between signatures were restricted to cells harboring antigen-reactive TCRs in the indicated class (melanocyte-reactive or viral-reactive).

#### 30,810 TCR library

Human paired TCR sequences were compiled from multiple sources including clinical trial NCT02305186^61,62^, and from whole blood and surgically resected tumor samples from patients with pancreatic ductal adenocarcinoma receiving treatment at Dana-Farber Cancer Institute (IRB #03-189 and #14-408). The study was conducted in accordance with the Declaration of Helsinki and the International Ethical Guidelines for Biomedical Research involving Human Subjects. Written informed consent was obtained from participants enrolled in #03-189 and #14-408 prior to sample collection.

For each PBMC sample, 6,000 cells were loaded into a 10X Chromium controller instrument along with Chromium Next GEM Single Cell 5’ v2 beads (10X Genomics PN-1000263). Up to four PBMC samples were multiplexed together after being tagged with unique DNA-barcoded antibodies as described above. For tumor samples, all sorted cells were loaded, and no multiplexing was performed. After RT-PCR, cDNA was purified, and a library was constructed from each sample using a 10X Library Construction Kit (10X Genomics PN-1000190) following the standard 10X protocol. An additional VDJ-enriched library was created for each sample using a specialized Chromium Single Cell Human TCR Amplification Kit (PN-1000252). Libraries were then sequenced on an Illumina NovaSeq system operated by Azenta/Genewiz generating paired end 150bp reads.

The following steps were performed separately for each patient. First, the TCR contigs sequenced in the pre- and post-treatment sample were collapsed into one list to allow for clonotype assignment independent of time. The list was filtered to eliminate any non-productive rearrangements and sequences obtained from cells that did not pass QC, as described above. Then, a tentative clonotype was assigned to each unique combination of 1 to 4 TCR chains that co-occurred in one cell; a TCR chain was defined as the productive combination of a V gene, D gene (for β chains only), J gene, and complementarity determining region 3 (CDR3). Clonotypes were finalized by reassigning any cell if its TCR chain set formed a subset of another clonotype. A clonotype was considered to be shared between the blood and the tumor sample of a given patient if at least one match occurred in the α-chain or the β-chain CDR3 amino acid sequence.

### Expression of TCRs in reporter Jurkat J76 cells

Jurkat J76 cells were lentivirally transduced to express CD8α and β chains in addition to a fluorescent reporter under the control of an nuclear factor of activated T cells (NFAT) response element^35^. Cells transduced by NFAT-CFP-encoding lentivirus were sorted as single cells into a 96-well plate and cultured until confluent. Half of each clonal population was stimulated with phorbol myristate acetate (PMA,10 ng/mL, Thermo Fisher) and ionomycin (1 ug/mL, Thermo Fisher). Reporter CFP expression was compared between stimulated cells and unstimulated cells and a clone with the highest differential was selected. The selected clone is henceforth referred to as NFAT-CFP CD8^+^ J76.

TCR-expressing lentivirus was produced by transiently transfecting HEK293T cells with psPAX2 packaging plasmid and pMD2.G VSV envelope plasmid, and TCR library transfer plasmid at a ratio of 5.6:3:1 (per T225 flask: 22.5 μg psPAX2, 7.5 μg pMD2.VSVG, 42 μg TCR library transfer plasmid) in addition to *Trans*IT-Lenti Transfection Reagent (Mirus Bio) at a 3:1 ratio of transfection reagent to DNA. Opti-MEM (Thermo Fisher), transfection reagent, and DNA were mixed and incubated for 10 minutes prior to dropwise addition to confluent HEK293T cells. Lentivirus was collected at 48 and 72 hours, centrifuged at 300*g* for 5 minutes to eliminate debris, and filtered through a 0.45-μm polyethersulfone filter (Millex, Millipore Sigma). Concentrated lentivirus (200×) was generated by ultracentrifugation at 100,000*g* for 45 minutes at 4C. Supernatant was discarded, and the pellet was resuspended overnight in 100 μL of Opti-MEM at 4C. Resuspended virus was aliquoted and stored at −80C.

Lentivirus was titered and NFAT-CFP CD8^+^ J76 cells were transduced at an MOI of 0.05. Transduced cells were sorted on a BD FACSAria at a minimum coverage of 1000× for the 3,808 TCR library and 100× for the 30,810 TCR library to generate TCR library-expressing reporter J76 cells.

For individual clonal TCR lines, TCR constructs were formatted as TCRβ-P2A-TCRα and cloned into the same pHIV backbone vector as the TCR library. NFAT-CFP CD8^+^ J76 cells were transduced with unconcentrated TCR lentivirus generated as described above (1 mL per 1M cells) and TCR expression was verified by flow cytometry with anti-TCR antibody (clone IP26, Biolegend).

### Library-versus-library RAPTR lentiviral screening assay

#### Assembly of RAPTR 101-pMHC virus library

Lentivirus activated by promoter shuffling (LeAPS) virus libraries were produced as previously described in Dobson et al.^28^. Briefly, HEK293T cells were seeded as described 24 hours pre-transfection. Libraries of 101 barcoded GFP-expressing pLeAPS plasmids and 101 pHIV-pMHC plasmids (Genscript) (**Table S3**) were used to generate VSVG-pseudotyped lentivirus in separate 24-well plates. Following virus collection at 48 hours, HEK293T cells were seeded at 20% confluency and transduced with pairs of barcoded LeAPS and pMHC viruses in duplicate 24-well plates (per well: 100 μL barcoded LeAPS virus and 700 μL pMHC virus) with the addition of 8 μg/mL of diethylaminoethyl-dextran (Sigma-Aldrich) to aid transduction.

After 48 hours, duplicate wells were examined via flow cytometry for transduction by pMHC virus (mCherry^+^) and LeAPS-barcode virus (GFP^+^). The duplicate wells in culture were pooled proportionally according to the proportion of pMHC- and LeAPS barcode-transduced (mCherry^+^ GFP^+^) cells to ensure proportional representation of each pMHC library member. Pooled cells were sorted (mCherry^+^ GFP^+^) to generate the 101-pMHC virus-packaging cell line. To generate pMHC virus, virus-packaging cells were seeded analogously as HEK293T cells for transfection with psPAX2.1 and pMD2.VSVG-mut plasmids using *Trans*It-Lenti (Mirus Bio) transfection reagent. 101-pMHC virus library was collected and concentrated at 48 and 72 hours as described above.

#### RAPTR viral library stimulation

To stimulate the 3,808 TCR library with the 101-pMHC virus library, 65 million cells were incubated in complete RPMI with 560 μL of concentrated 101-pMHC virus library and 8 μg/mL of diethylaminoethyl-dextran (Sigma-Aldrich) for 24 h at 37C. Cells were washed once in FACS buffer and NFAT-CFP^+^ cells were sorted on a BD FACSAria cell sorter.

#### RAPTR scRNA-seq of transduced cells

Prior to single-cell sequencing cells to extract TCRs enriched by stimulation with the 101-pMHC virus library, cells were sorted for transduction (GFP^+^). Cells were analyzed using the 10X Genomics Chromium GEM-X Single Cell 5’ v3 kit (PN-1000699). 0.5 μL of 10 μM custom TCR-specific primer (10X_TRAC_RT) was spiked into the reverse transcription (RT) mix to maximize TCR capture and a pMHC barcode-construct specific primer was added to the cDNA amplification mix (1 μL of 10 μM 10X_pMHC_cDNA) followed by a 0.65× SPRIselect bead (Beckman Coulter) cleanup. TCR amplicons were generated from cDNA via two nested PCRs: PCR #1 (98C for 45 sec; 15 cycles of 98C for 20 sec, 62C for 30 sec, 72C for 30 sec; 72C for 1 min) with 0.2 μM of 10X_nested_f and 10X_TCR_outer_r using KAPA HiFi HotStart ReadyMix (Roche) followed by a left-sided 0.8× SPRIselect bead cleanup; PCR #2 (98C for 45 sec; 13 cycles of 98C for 20 sec, 62C for 30 sec, 72C for 30 sec; 72C for 1 min) with 0.2 μM of 10X_nested_f and 10X_TCR_inner_r followed by a left-sided 0.8× SPRIselect bead cleanup. pMHC amplicons were generated from cDNA via PCR (98C for 45 sec; 25 cycles of 98C for 20 sec, 62C for 30 sec, 72C for 30 sec; 72C for 1 min) with 0.2 μM of 10X_nested_f and 10X_pMHC_r followed by a left-sided 0.8× SPRIselect bead cleanup. TCR and pMHC amplicons were indexed, pooled, and sequenced (150 bp PE) on an Element AVITI.

#### RAPTR scRNA-seq analysis

Cell-feature matrices were constructed for both TCR and pMHC amplicons using 10X CellRanger (v9.0.1). Unique 44-bp CDR3β barcodes were used as feature references to identify TCRs and 18-bp barcodes were used to identify pMHCs. Downstream analysis was completed in Python. In each dataset, cells with < 5 TCR or pMHC UMIs and cells with less than 60% of UMIs corresponding to a single TCR identity were filtered out. Each cell was assigned a TCR or pMHC identity based on the highest number of UMIs for each. The TCR and pMHC datasets were merged on cell barcodes, ensuring each cell had both a TCR and pMHC identity. TCR identities with fewer than 5 cells were filtered out (46 of 3,202 cells). Plots were generated using Matplotlib and Seaborn packages.

### Peptide-APC and J76 library co-culture screen

Peptides were selected from several sources for screening using antigen presenting cells. The Immune Epitope Database (IEDB) was used to assemble a list of HLA-A2-restricted human self-epitopes derived from genes that had been previously confirmed as immunogenic in T cell assays in the context of vitiligo or melanoma. To identify novel candidate antigens specific to melanocytes, scRNA-seq data from melanocytes isolated from vitiligo donors were integrated with melanocyte scRNA-seq data from 17 donors profiled in Gellatly et al^50^. Genes specifically expressed in melanocytes were identified by performing differential expression analysis between melanocytes and all other cell types. Genes significantly upregulated (FDR-adjusted *P* < 0.05) and exhibiting a log fold change > 1 were selected. This set was then intersected with melanocyte-specific gene lists derived from external datasets.

Melanocyte-specific gene expression in external datasets was identified using a multi-step data integration approach leveraging three datasets: the FANTOM5 project^76^, GTEx Portal^77^, and the single-cell analysis study by Belote et al^78^. In the FANTOM5 database, genes were selected if their expression was more than twofold higher than in any other tissue and exceeded 5 Transcripts Per Million (TPM) in the melanocyte categories of “Melanocyte.dark,” “Melanocyte.light,” or “Melanocyte.” This yielded 227 candidate genes. Gene expression data from the GTEx Portal were then used to further refine the list to focus on skin-specific expression. Genes with higher expression in “Skin – Sun Exposed (Lower leg)” and “Skin – Not Sun Exposed (Suprapubic)” compared to all other non-brain and non-nerve tissues were prioritized, resulting in 13 genes with elevated expression in these skin tissues. Expression of proteins from these genes in melanocytes was then confirmed using a dataset from Belote et al.^78^, leading to the identification of five known melanocyte-specific genes: *PMEL, MLANA, TYRP1,* and *DCT*. Excluding these genes that have been extensively examined in vitiligo, 10 genes were selected for screening (*CD63, CDH3, CYGB, GMPR, GPR143, PLP1, SLC1A4, SOX10, VAT1*). Epitopes predicted to bind HLA-A2 were selected for screening using NetMHCpan version 4.1. Epitopes associated with extracted TCRs in VDJDb^59^ were also included. We further assembled a list of melanoma-associated epitopes from literature^37,39,40^.

All peptides (Genscript) were pooled as described in **Table S3** at 5 mg/mL in dimethyl sulfoxide. TAP-deficient T2 cells were pulsed with peptide pools at 10 μg/mL or individual peptides at 1 μg/mL for 4 h and incubated at a 1:1 ratio with TCR-expressing NFAT-CFP CD8^+^ J76 cells at 37C for 20-24 h. Individual peptide validation was completed in 96-well U-bottom plates. Cells were washed and stained with anti-CD69 (Biolegend, clone FN50) and anti-CD19 (Biolegend, clone HIB19) in FACS buffer, both at a 1:200 dilution. Cells were analyzed for activation (NFAT^+^CD69^+^) on a Cytoflex S flow cytometer. For peptide pools, activated cells were sorted with a BD FACSAria cell sorter.

### Bulk sequencing of TCRs enriched by antigen screens

To identify TCRs enriched by screens with the 101-pMHC virus library or peptide-pulsed T2 cells, genomic DNA was isolated from J76 cells using the PureLink Genomic DNA kit (Thermo Fisher). Noting that CDR3α and CDR3β were ordered on a single oligo and that each was codon-optimized to uniquely encodes a specific TCR, extracting one of the CDR3α or CDR3β enabled identification of the full TCR clonotype. Using the CDR3β for identifying TCRs in this assay, TRBV-CDR3β amplicons were amplified (98C for 30 sec; 24 cycles of 98C for 30 sec, 70C for 30 sec, 72C for 45 sec; 72C for 1 min) from 1 μg of genomic DNA using NEBNext Ultra II Q5 Master Mix (NEB) and 1 μM of each forward and reverse primer (for TCR: TRBV_CDR3B_f and TRBV_CDR3B_r; for pMHC: pMHC_BC_f and pMHC_BC_r). Amplicons were submitted for Amplicon-EZ analysis by Genewiz. Enrichment was calculated for each TCR as the fraction of reads for each enriched TCR divided by the TCR frequency in the base TCR library cell line quantified by sequencing on an Element AVITI (300 bp PE).

### Individual functional TCR validation

Monoclonal TCR lines were established by assembling individual TCRs in pHIV backbones, generating unconcentrated TCR lentivirus as described above and transducing 1 million NFAT-CFP CD8 J76 cells with 1 mL of unconcentrated virus. T2 cells were pulsed with individual peptides (Genscript) at 1 μg/mL for 2-4 hours. 100,000 peptide-pulsed T2 cells were incubated with 100,000 monoclonal TCR-expressing NFAT-CFP CD8 J76 cells overnight. Cells were washed and stained with anti-CD69 (Biolegend, clone FN50) and anti-CD19 (Biolegend, clone HIB19) in FACS buffer (PBS + 0.1% BSA + 1 mM EDTA) (1:200 dilution), before analysis on a Cytoflex S flow cytometer.

#### Antibodies in flow cytometry

All antibodies were used at a 1:50 or 1:200 dilution from stock concentration as described. Cells were stained in FACS buffer (PBS + 0.1% BSA + 1 mM EDTA) for 20 minutes at 4C, washed, and sorted on a BD FACSAria or analyzed on a Cytoflex S. All antibodies are from BioLegend.

#### Quantification and statistical analyses

Statistical analyses were performed using Python or GraphPad Prism (v.10). Information on specific statistical tests is included in figure legends. Data in bar plots is represented as the mean ± S.D. as indicated in figure legends. P values are listed in figure legends.

#### Software

Graphs were generated using Python and GraphPad Prism (v.10). Flow cytometry data were analyzed by FlowJo (v.10.10.0).

## RESOURCE AVAILABILITY

### Lead contact

Further information and requests for resources and reagents should be directed to and will be fulfilled by the lead contact, Michael Birnbaum.

### Materials availability

TCRAFT plasmids will be available as kits on Addgene (Birnbaum group) before publication. Additional plasmids are available upon request.

### Data and code availability

Next-generation sequencing and RAPTR scRNA-seq datasets are available in the National Center for Biotechnology Information Sequence Read Archive under accession number PRJNA1247142. Code to generate oligo pools for TCRAFT, analyze library composition, and process NGS and single-cell sequencing data is available on Github at https://github.com/birnbaumlab/TCRAFT/ and https://github.com/birnbaumlab/Gaglione-et-al-2025. Any additional information required to reanalyze the data reported in this paper is available from the lead contact upon request.

## Acknowledgements

We thank the Koch Institute’s Robert A. Swanson (1969) Biotechnology Center for their technical support, especially the Flow Cytometry Facility and MIT BioMicro Center. We thank S. Levine, N. Kamelamela, and G. Paradis for helpful discussions and suggestions. This work was supported in part by the Koch Institute Frontier Research Program through the Michael (1957) and Inara Erdei Fund and the Casey and Family Foundation Research Fund, the Packard Foundation, NIH Director’s New Innovator Award (DP2-AI158126), U.S. Army Medical Research (W81XWH2210300), and Pfizer Inc. to M.E.B.; Hartford Foundation, Vitiligo Research Fund, NIH AATG T32 (AI132152), and NIH P50 (AR080593-01) to J.E.H.; a Canadian Institutes for Health Research Doctoral Foreign Study award to S.A.G.; a Medical Scientist Training Program grant (T32 GM007753) from the National Institute of General Medical Sciences to B.E.S.; a National Science Foundation Graduate Research Fellowship and fellowship from Ludwig Center at MIT’s Koch Institute to C.R.P; a graduate research fellowship from the Ludwig Center at MIT’s Koch Institute to E.J.K.X. This work was delivered as part of the MATCHMAKERS team, of which M.E.B. is a member, supported by the Cancer Grand Challenges partnership financed by CRUK (CGCATF-2023/100001), the National Cancer Institute (OT2CA297463), and The Mark Foundation for Cancer Research. S.K.D. and H.S. are supported by the Hale Center for Pancreatic Cancer Research at Dana-Farber Cancer Institute (DFCI). S.K.D is a member of the Parker Institute for Cancer Immunotherapy at DFCI. M.E.B. and S.K.D. are supported by Break Through Cancer. M.E.B. and M.D. are supported by the Bridge Project, a partnership between the Koch Institute for Integrative Cancer Research at MIT and the Dana-Farber/Harvard Cancer Center. This work was additionally supported in part by the Koch Institute Support (core) Grant P30-CA14051 from the National Cancer Institute. Vitiligo samples were obtained from subjects who provided written consent to be included in Protocol H-14848. We would like to thank all our subjects who agreed to participate in this study. This content is solely the responsibility of the authors and does not necessarily represent the official views of the National Institutes of Health or National Cancer Institute.

## Author contributions

Conceptualization, S.A.G., R.S.M., and M.E.B; methodology, all authors; data analysis, S.A.G., R.S.M., C.K., M.H.W., S.A.J., L.R.A., J.S., and P.V.H.; sample acquisition, E.L.K., K.J.G., L.R.A., J.S., H.S., M.G., M.D., S.K.D., and J.E.H.; writing, S.A.G., C.K., and M.E.B.; review and editing, all authors.

