## Supplementary Information for "Scalable TCR synthesis and screening enables antigen reactivity mapping in vitiligo"

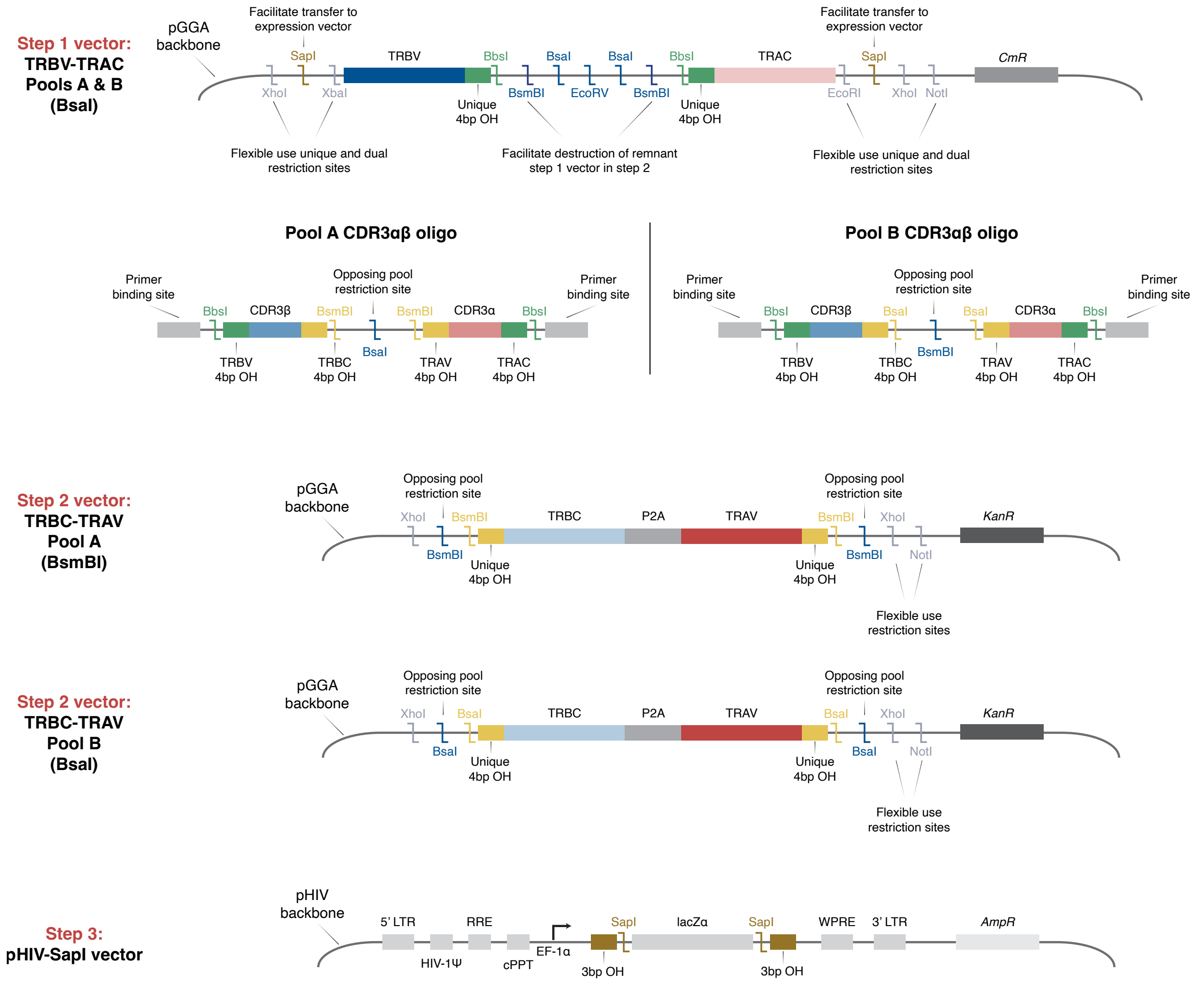


**Supplementary Figure 1. Design of TCRAFT vectors (TRBV-TRAC, TRBC-TRAV, destination vector) and oligo pools**. Schematic showing components and enzyme sites of designed vectors and oligo pools for seamless and efficient pairing of TCR components. TRAV and TRBV alleles were split into two pools each by design to maximize assembly efficiency.

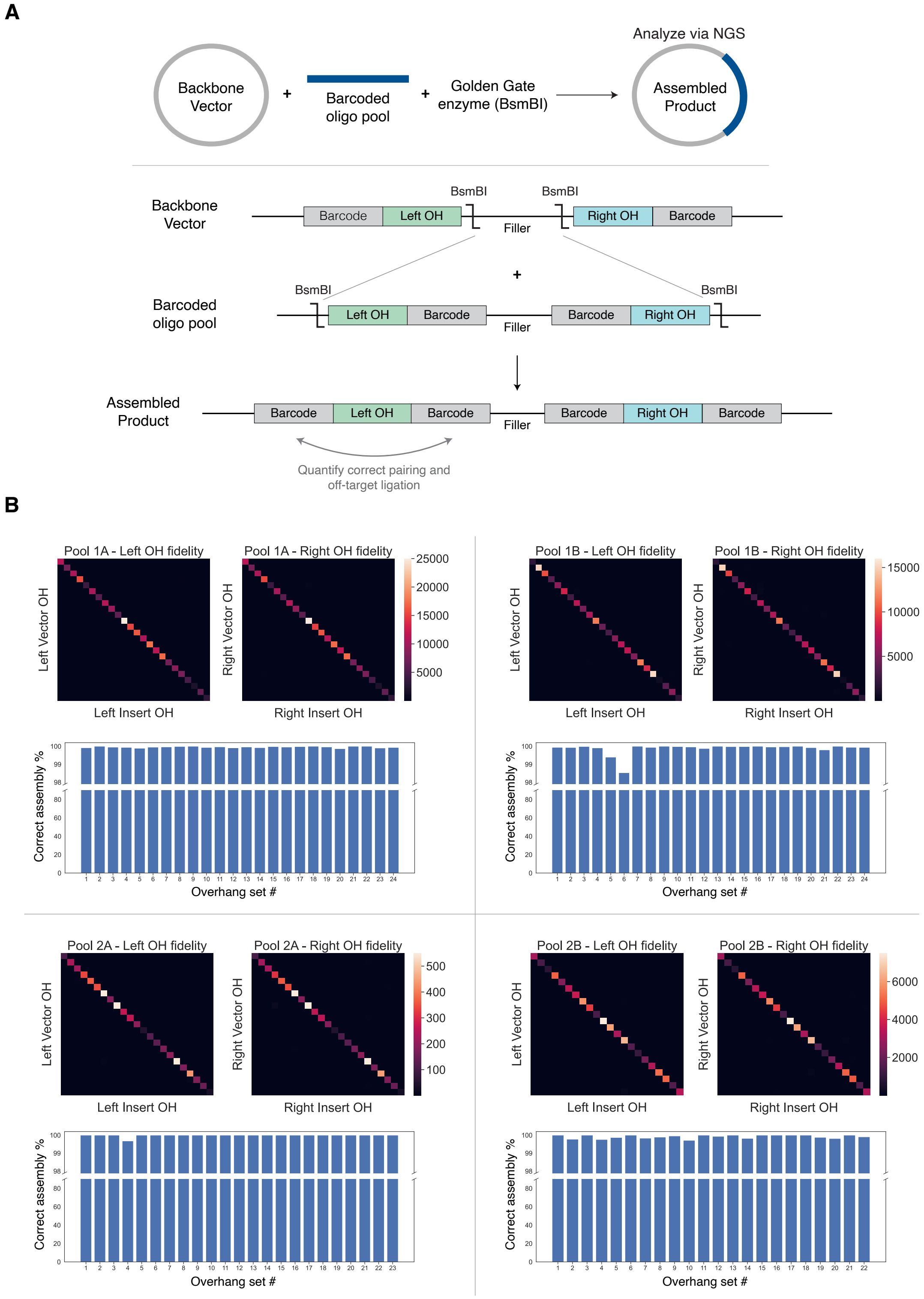


**Supplementary Figure 2. Experimental validation of overhang sets. (A)** Schematic overview of overhang validation experiment. Four pools of backbone vectors containing overhang sets and barcodes were mixed with corresponding oligo pools. Comparison of barcodes on either side of the barcode allows for quantification of on-target and off-target vector-insert pairing. **(B)** Fidelity of left and right overhangs and overall assembly efficiency of each designed overhang set.

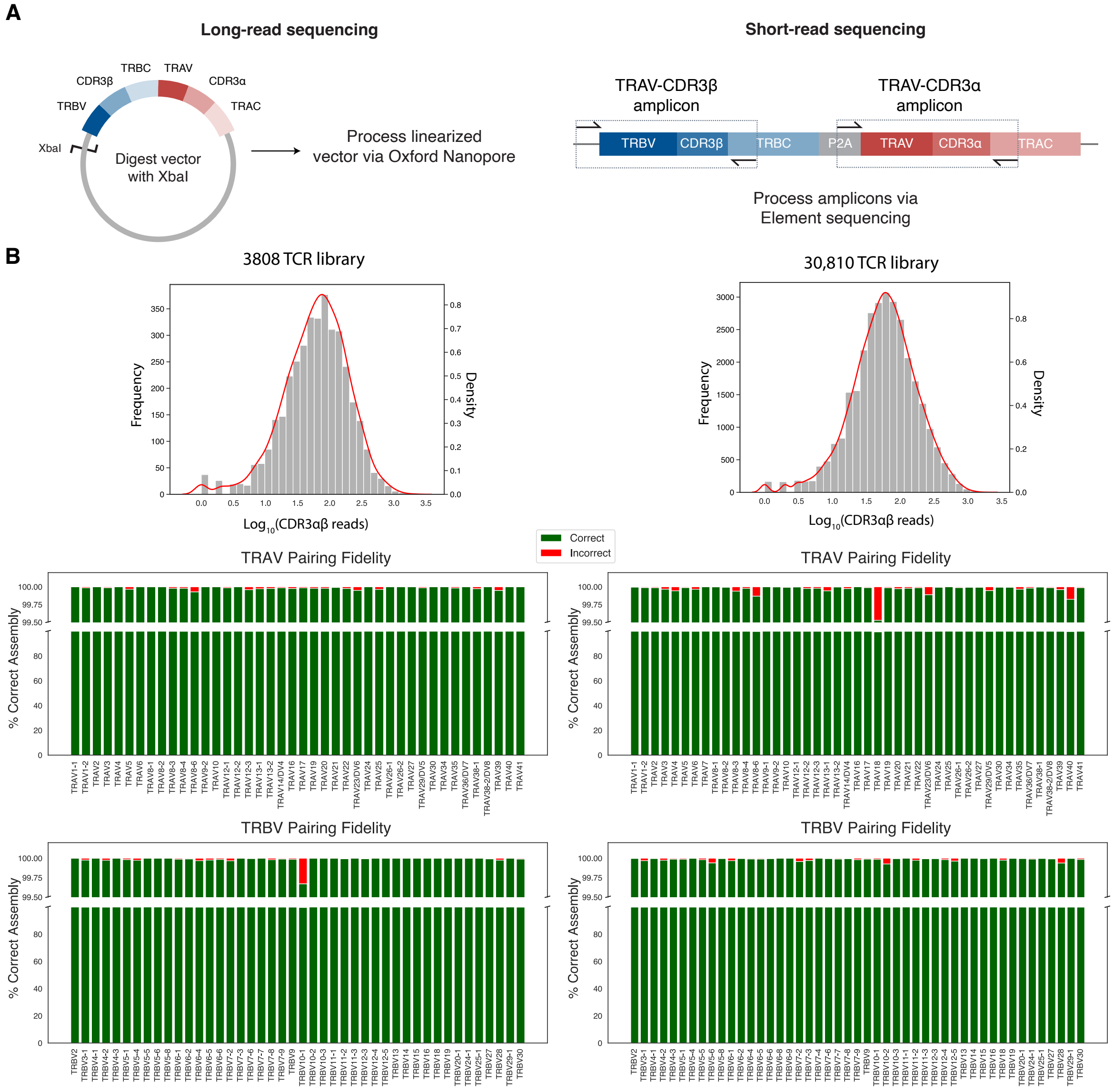


**C**

| Library | Step | CFU/uL | Electroporations | Coverage |
|---|---|---|---|---|
| 3,808 TCR | 1A | 1.24e7 | 3 | 4880x |
| | 1B | 3.6e6 | 3 | 1420x |
| | 2A | 5.7e6 | 6 | 8980x |
| | 2B | 4.6e7 | 7 | 8450x |
| | 3 | 3.8e6 | 8 | 8060x |
| 30,810 TCR | 1A | 6.8e6 | 15 | 7230x |
| | 1B | 4.0e6 | 15 | 3590x |
| | 2A | 3.8e6 | 14 | 1860x |
| | 2B | 2.4e6 | 15 | 1160x |
| | 3 | 1.85e6 | 20 | 1200x |

**Supplementary Figure 3. TCR assembly sequencing and efficiency. (A)** Vector and amplicon preparation for sequencing. Long-read sequencing: vectors are digested with a single restriction enzyme prior to ONT sequencing. Short-read sequencing: TRBV-CDR3β and TRAV-CDR3α regions are PCR amplified and sequenced via Element AVITI. **(B)** Frequencies of complete, correctly assembled TCR sequences and proportion of TCRs correctly assembled for each TRAV and TRBV allele for 3,808 and 30,810 TCR libraries (ONT). **(C)** Electroporation efficiency per μL of reaction product at each step (electrocompetent DH10B *E. coli*), number of electroporation transformations per step, and coverage per TCR.

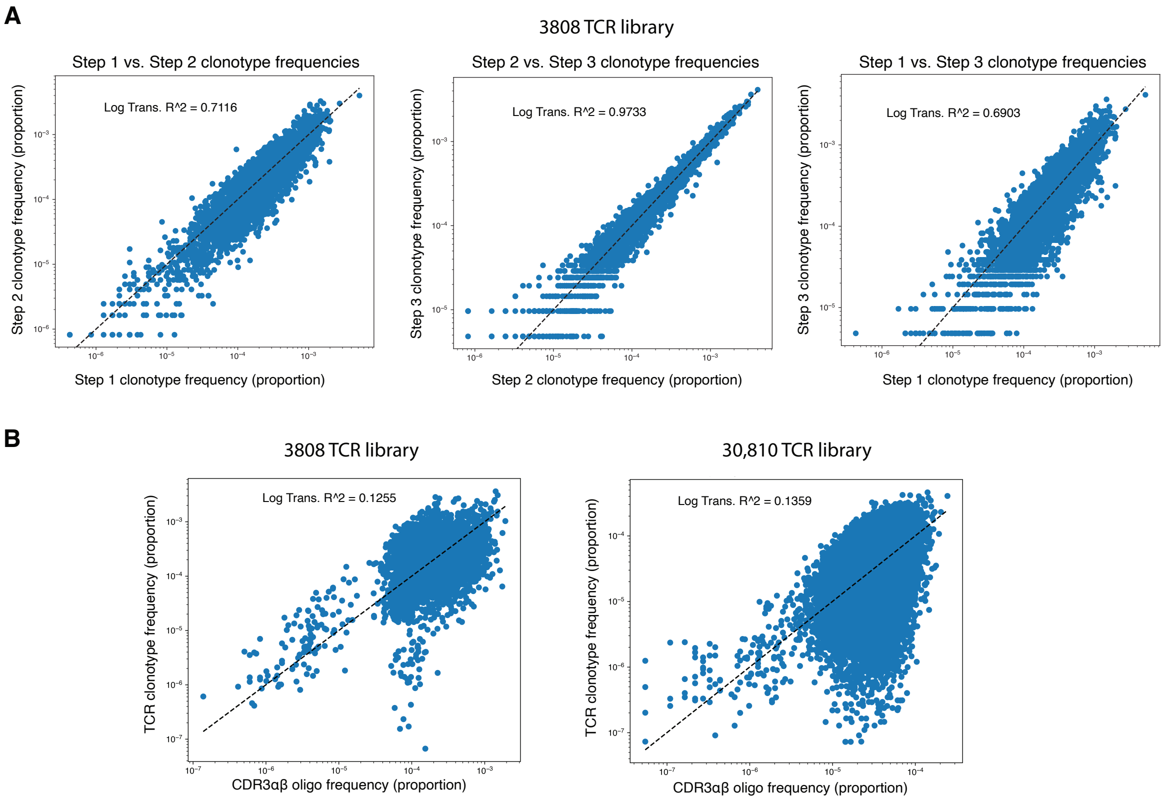


**Supplementary Figure 4. 3808 TCR clonotype and oligo frequencies across TCRAFT workflow. (A)** Correlations between frequencies of clonotypes across reaction steps (step 1 = oligos inserted into TRBV-TRAC vectors, step 2 = TRBC-TRAV inserted into TRBV-CDR3β-CDR3α-TRAC to form complete TCR library in pGGA cloning vector, step 3 = complete TCR library in destination pHIV expression vector). **(B)** Oligo frequency versus TCR frequency (average of CDR3α and CDR3β frequency).

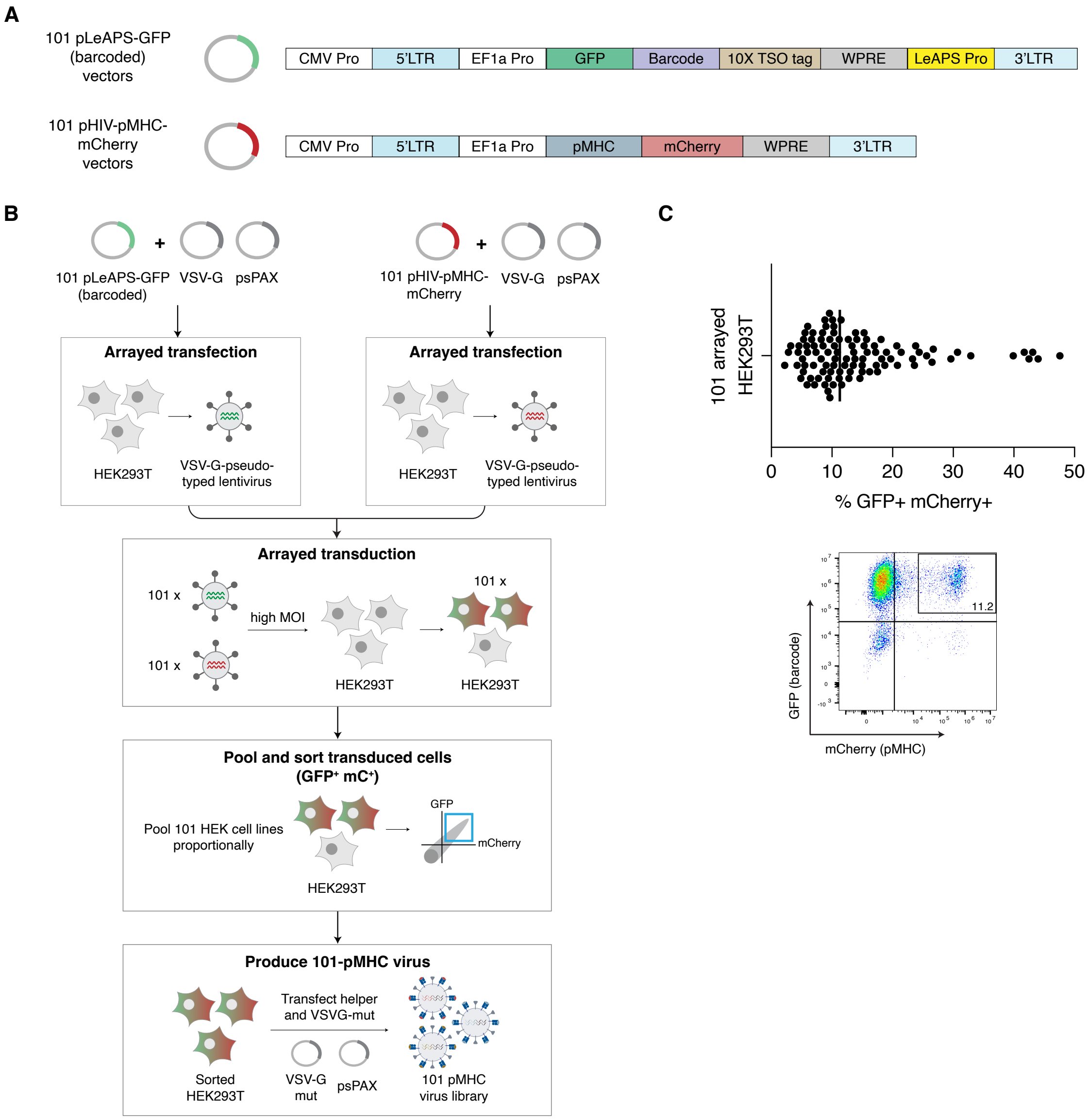


**Supplementary Figure 5. Generation of 101-pMHC virus library with pLeAPS. (A)** Format of 101 pLeAPS-GFP barcode and pHIV-pMHC-mCherry lentiviral vectors used to construct packaging cells for the 101-pMHC virus library. Each pLeAPS-GFP barcode vector includes a fluorescent marker (GFP), a 10X capture tag corresponding to the 10X TSO sequence[36], an 18-bp barcode, and the LeAPS CMV promoter[28]. Each pHIV-pMHC-mCherry vector encodes a pMHC in single-chain trimer format. **(B)** Strategy for constructing 101-pMHC virus library packaging cell line. 101 pLeAPS-GFP (distinct barcodes) and 101 pMHC-pMHC vectors are separately transfected in arrayed format with helper plasmids to generate VSV-G-pseudotyped lentiviruses. HEK293T cells are transduced in array, each well receiving one barcode LeAPS-VSVG virus and one pMHC-VSVG virus (101 wells total). The pHIV-pMHC-mCherry virus drives pMHC expression on the surface of HEK293T cells, while the LeAPS virus delivers a repackageable genome encoding a fluorophore and barcode. The 101 transduced cell populations are pooled proportionally and sorted for dual expression of GFP and mCherry (GFP$^{+}$mC$^{+}$) in a single sorting step. Sorted cells are then transfected with helper plasmids (psPAX, VSV-G-mut) to generate the 101-pMHC virus library. This packaging cell line can be maintained and reused indefinitely for virus library production via transfection with helper plasmids. (**C)** Arrayed transduction of HEK293T cells with 101 pMHC and LeAPS viruses (GFP$^{+}$mC$^{+}$), prior to proportional pooling and sorting. Representative flow plot of arrayed pMHC-LeAPS dual transduction.

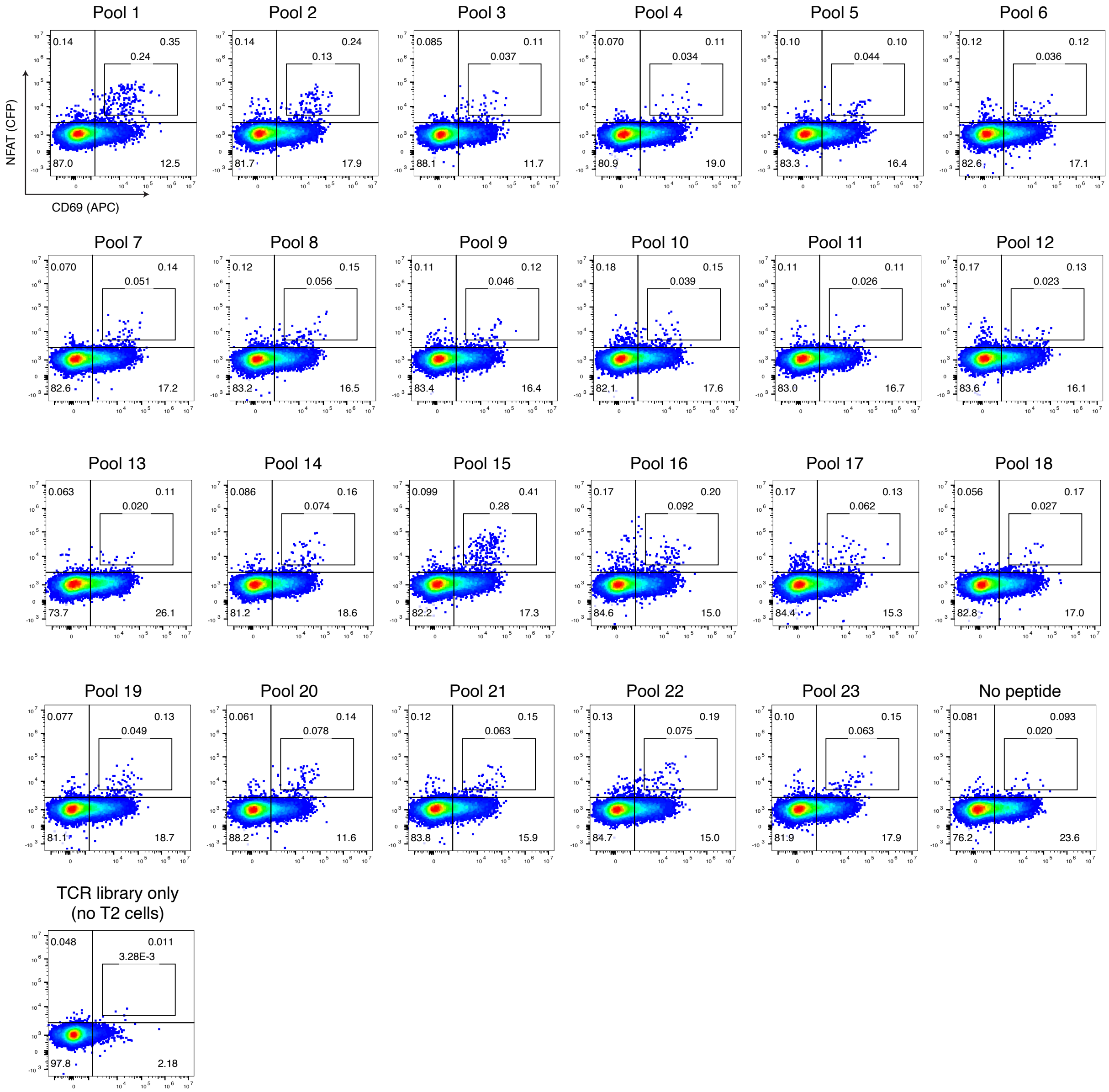


**Supplementary Figure 6. Peptide pool activation of 3,808 TCRs.** Representative flow plots for 23 peptide pools corresponding to Figure 4.

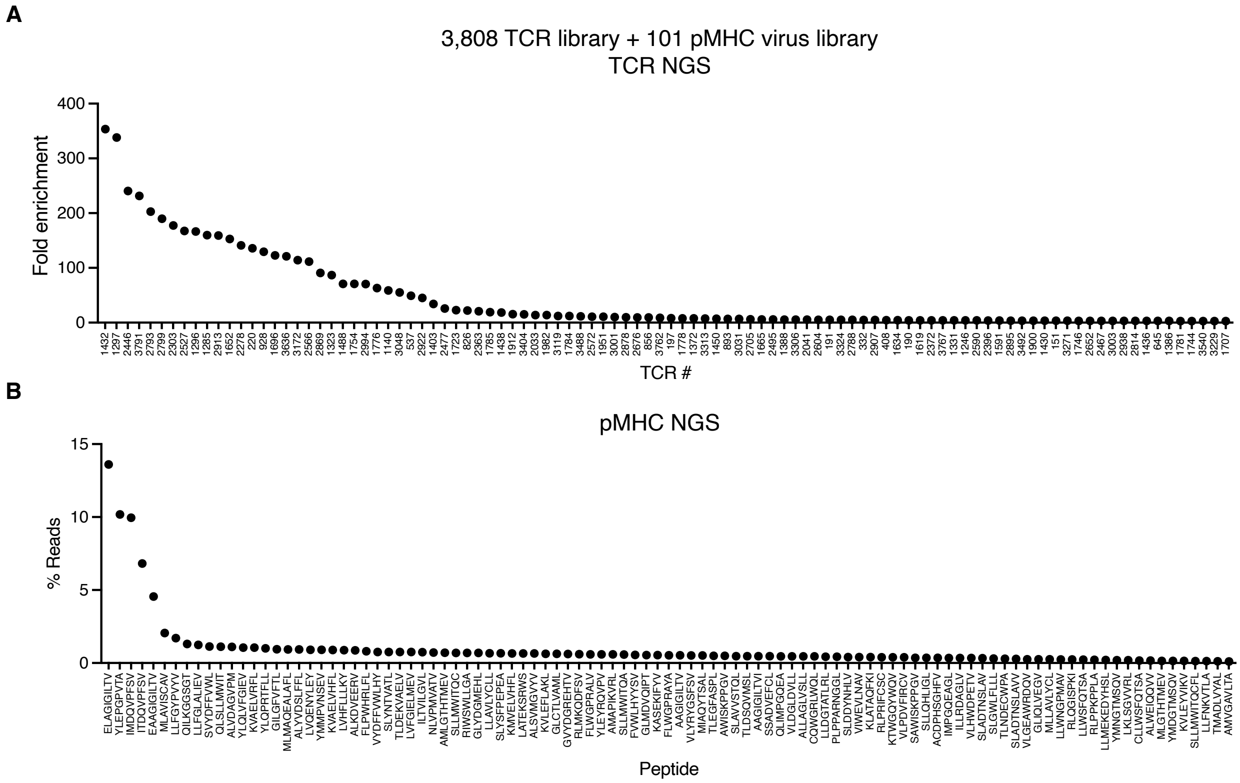


**Supplementary Figure 7. RAPTR captures reactive TCRs and antigens via NGS. (A)** Fold enrichment of TCRs in cells activated by 101-pMHC virus library compared with naïve 3,808 TCR library. **(B)** Proportion of barcode (pMHC) reads in cells activated and transduced by 101-pMHC virus library.

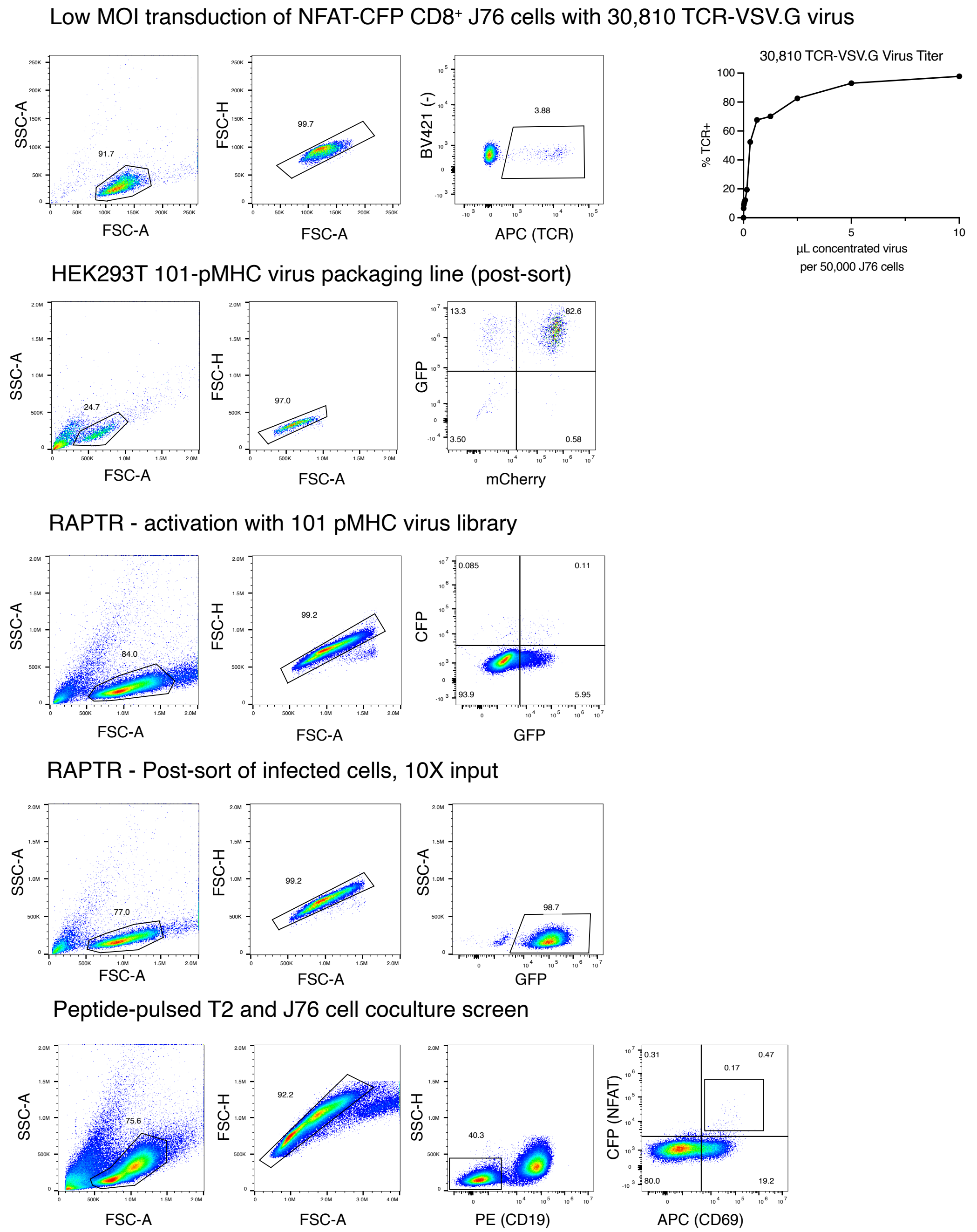


**Supplementary Figure 8. Gating strategies.** Flow cytometry gating strategies used throughout manuscript.

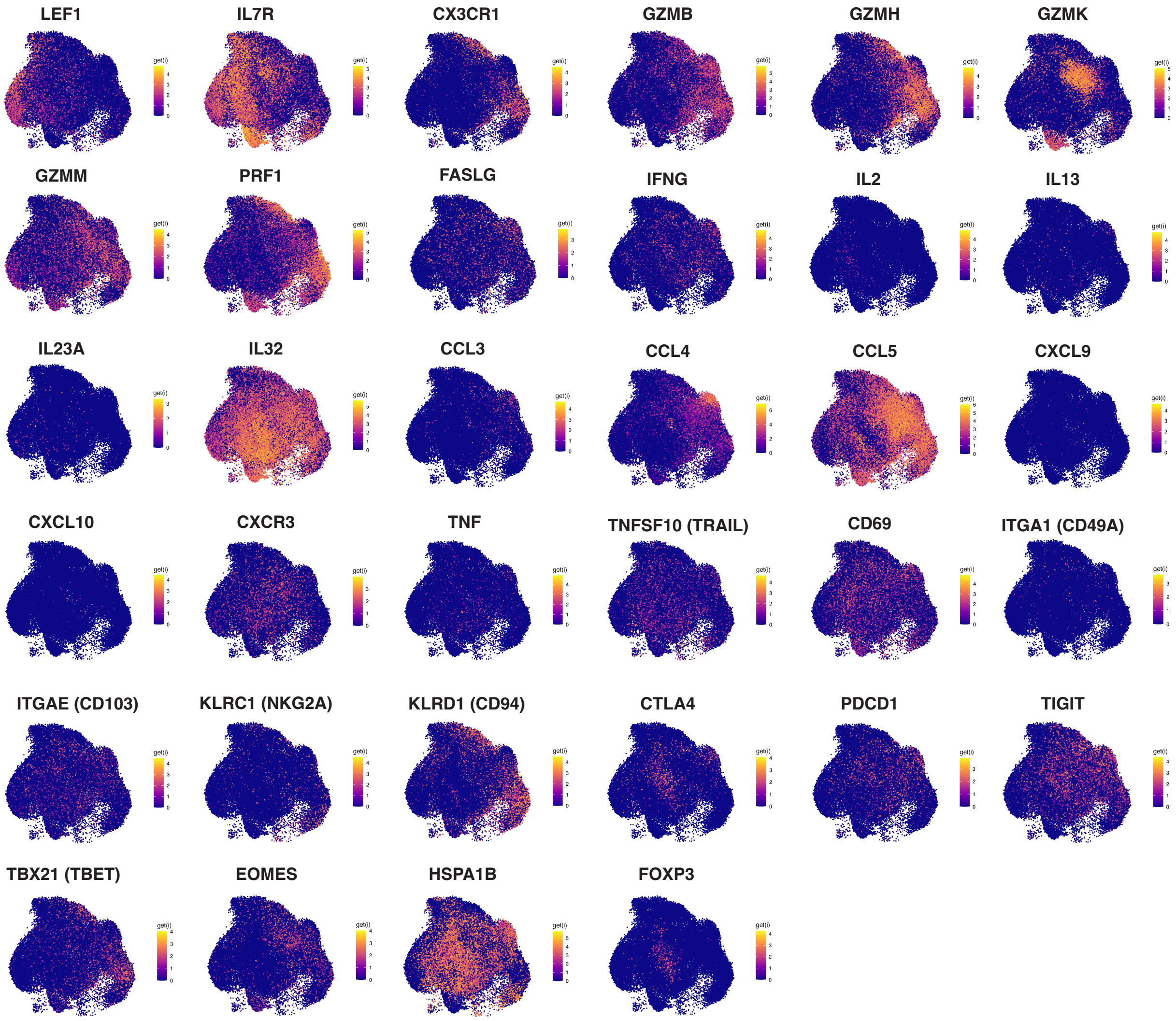


**Supplementary Figure 9**. Expression of additional markers across all cells in vitiligo T cell atlas presented in Figure 5A.

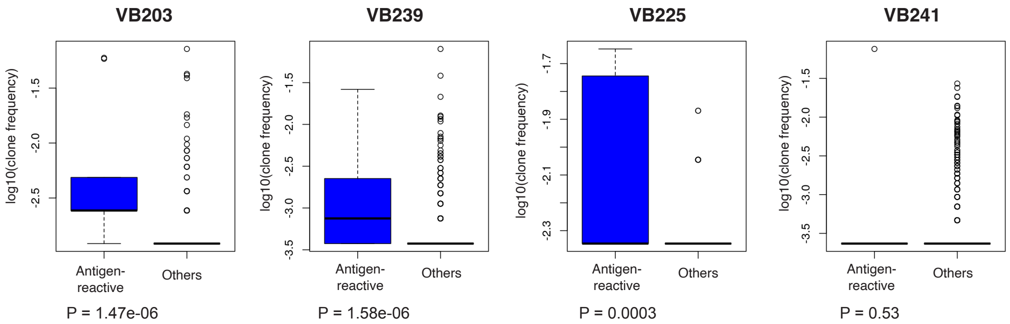


**Supplementary Figure 10. Clonal frequency distribution for antigen-detected clones vs. non-antigen detected clones (others).** Data are shown for donors (VB203, VB239, VB225, VB241) with at least 5 melanocyte-reactive TCRs. The full list of antigen-reactive TCRs for all donors is shown in Table S7.

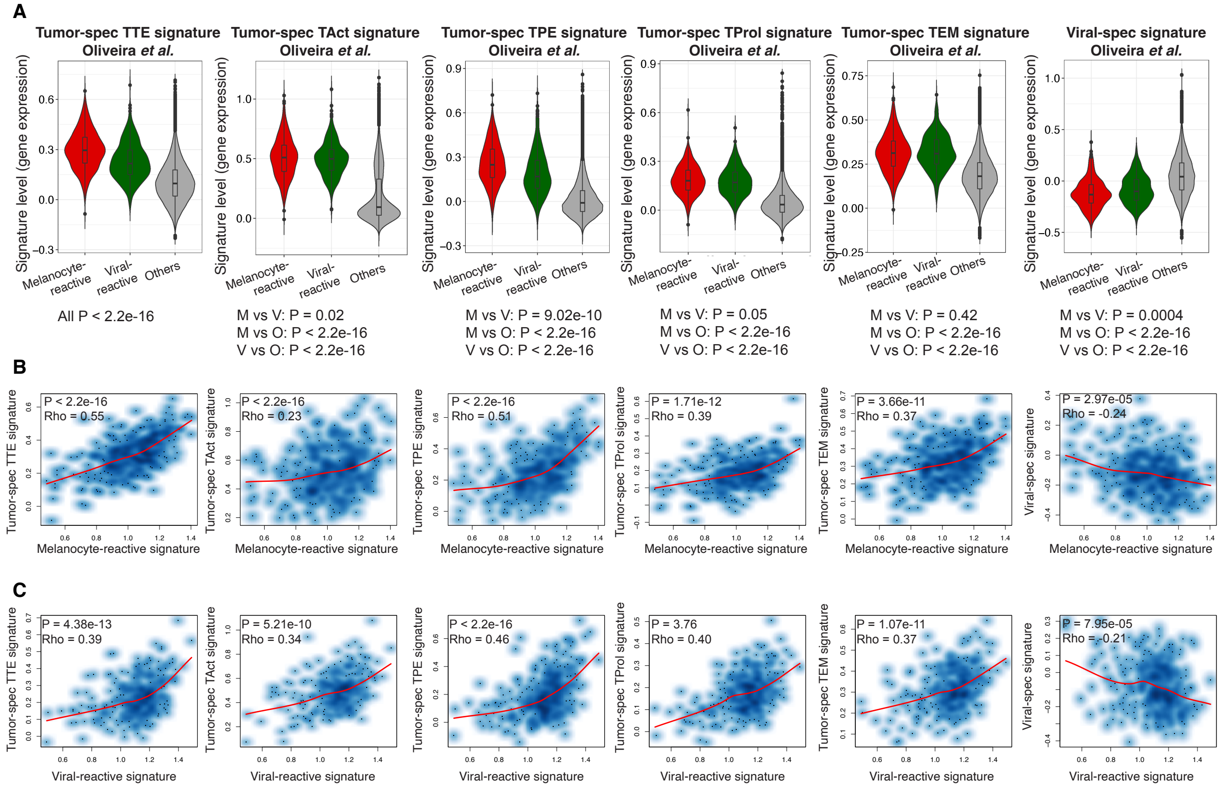


**Supplementary Figure 11. Gene signatures of melanocyte-reactive, viral-reactive, and melanoma antigen-reactive TCRs from Oliveira et al.**[7] **(A)** Application of melanoma antigen-reactive and viral antigen-reactive TCR gene signatures from Oliveira et al.[7] to clonotypes in our dataset. **(B)** Correlation of the melanocyte-reactive gene expression signature with gene signatures of melanoma antigen-reactive or viral antigen-reactive TCRs across all cells harboring antigen-reactive TCRs in our dataset. **(C)** Correlation of the viral-reactive gene expression signature with gene signatures of melanoma antigen-reactive or viral antigen-reactive TCRs across all cells expressing antigen-reactive TCRs in our dataset.
